# Regulation of metabolic health by dietary histidine in mice

**DOI:** 10.1101/2022.04.24.489217

**Authors:** Victoria Flores, Alexandra B. Spicer, Michelle Sonsalla, Nicole E. Richardson, Deyang Yu, Grace E. Sheridan, Michaela E. Trautman, Reji Babygirija, Eunhae P. Cheng, Jennifer M. Rojas, Shany E. Yang, Matthew H. Wakai, Ryan Hubbell, Ildiko Kasza, Jay L. Tomasiewicz, Cara L. Green, Claudia Dantoin, Caroline M. Alexander, Joseph A. Baur, Kristen C. Malecki, Dudley W. Lamming

**Affiliations:** William S. Middleton Memorial Veterans Hospital, Madison, WI 53705, USA; Department of Medicine, University of Wisconsin-Madison, Madison, WI 53705, USA; Interdepartmental Graduate Program in Nutritional Sciences, University of Wisconsin-Madison, Madison, WI 53706, USA; Department of Population Health Sciences, School of Medicine and Public Health, University of Wisconsin-Madison, Madison, WI 53726, USA; Endocrinology and Reproductive Physiology Graduate Training Program, University of Wisconsin-Madison, Madison, WI 53706, USA; Molecular and Environmental Toxicology Program, University of Wisconsin-Madison, Madison, WI 53706, USA; Graduate Program in Cellular and Molecular Biology, University of Wisconsin-Madison, Madison, WI 53706, USA; Department of Physiology and Institute for Diabetes, Obesity, and Metabolism, University of Pennsylvania, Philadelphia, PA 19104, USA; McArdle Laboratory for Cancer Research, University of Wisconsin-Madison, Madison, WI 53705, USA; University of Wisconsin Carbone Cancer Center, Madison, WI 53705, USA

**Author notes:** Correspondence and Lead ContactDudley W. Lamming, PhD Associate Professor of Medicine University of Wisconsin-Madison 2500 Overlook Terrace, VAH C3127 Research 151 Madison, WI 53705, USA.

**Keywords:** histidine, amino acids, protein restriction, BMI, body composition, FGF21

## Abstract

Low protein (LP) diets are associated with a decreased risk of diabetes in humans, and a low protein diet promotes leanness and glycemic control in both rodents and humans. While the effects of a LP diet on glycemic control are mediated by reduced dietary levels of the branched-chain amino acids (BCAAs), we have observed that reducing dietary levels of the other six essential amino acids leads to changes in body composition. Here, we find that dietary histidine plays a key role in the response to a LP diet in male C57BL/6J mice. Specifically reducing dietary levels of histidine by 67% reduces weight gain of young, lean male mice, reducing both adipose and lean mass gain, without altering glucose metabolism. Specifically reducing dietary histidine rapidly reverses diet-induced obesity and hepatic steatosis in diet-induced obese male mice, increasing insulin sensitivity; this normalization of metabolic health was associated not with caloric restriction or increased activity, but with increased energy expenditure. We find that the effects of histidine restriction surprisingly does not require the energy balance hormone *Fgf21*. Histidine restriction started in mid-life promoted leanness and glucose tolerance in aged males but not females, but did not affect frailty or lifespan in either sex. Finally, we demonstrate that variation in dietary histidine levels helps to explain body mass index differences in humans. Overall, our findings demonstrate that dietary histidine is a key regulator of weight and body composition in male mice and in humans, and suggest that reducing dietary levels of histidine may be a highly translatable option for the treatment of obesity.

**Key Points:**

- Protein restriction (PR) promotes metabolic health in rodents and humans and extends rodent lifespan.
- Restriction of specific individual essential amino acids can recapitulate the benefits of PR.
- Reduced histidine promotes leanness and increased energy expenditure in mice.
- Reduced histidine does not extend the lifespan of mice when begun in mid-life.
- Dietary levels of histidine are positively associated with BMI in humans.

## Introduction

Over the last two decades, it has become clear that calories from different macronutrient groups differ in their ultimate metabolic impact and effects on health and longevity, and that the traditional view of food and obesity – that a calorie is “just a calorie” – is too simple. Multiple studies have found that calories from carbohydrates, fat, and protein all have unique effects on human metabolism (Hall *et al*., 2015; Fontana *et al*., 2016). Calories from protein have generally been thought of as beneficial, as dietary protein promotes satiety and displaces carbohydrates and calorie-dense fat; further, some metabolic benefits of dietary protein have been noted in short-term studies (Seino *et al*., 1983; Gannon *et al*., 2003; Dong *et al*., 2013). However, multiple long-term observational studies have found that humans eating a low protein (LP) diet have a reduced incidence of diabetes and other age-related diseases (Linn *et al*., 2000; Lagiou *et al*., 2007; Sluijs *et al*., 2010; Levine *et al*., 2014; van Nielen *et al*., 2014). In a short-term randomized clinical study, an LP diet promotes leanness and glycemic control in humans with obesity (Fontana *et al*., 2016). LP diets also improve metabolic health in rodents, promoting leanness and blood glucose control in the context of both lean and diet-induced obese animals, and increasing lifespan (Laeger *et al*., 2014; Solon-Biet *et al*., 2014; Solon-Biet *et al*., 2015; Fontana *et al*., 2016; Maida *et al*., 2016; Cummings *et al*., 2018; Richardson *et al*., 2021).

We hypothesized that that the beneficial effects of an LP diet might be the results of reduced consumption of specific essential amino acids. In initial studies conducted over the last decade, we focused on the branched-chain amino acids (BCAAs; leucine, isoleucine, and valine), which have been linked to insulin-resistant obesity and diabetes in both rodents and humans. We demonstrated that a 67% restriction of all three BCAAs (or isoleucine alone) recapitulates the effects of an LP diet in lean and Western diet-induced obese mice, enhancing leanness and blood glucose control; we have also observed beneficial effects of an LP diet on aged wild-type mice (Fontana *et al*., 2016; Cummings *et al*., 2018; Richardson *et al*., 2021; Yu *et al*., 2021). Intriguingly though, we observed that a diet with a 67% restriction of the other six essential amino acids – histidine, lysine, methionine, phenylalanine, threonine, and tryptophan – resulted in reduced weight gain and adiposity without improving glucose tolerance (Fontana *et al*., 2016).

Here, we investigate the role of these other six essential amino acids, examining the effects of reducing each of these amino acids individually. We find that that restriction of either histidine, phenylalanine, or threonine is sufficient to reduce adiposity in young mice, but does not alter blood glucose control. As restriction of dietary threonine has recently been shown to improve metabolic health by another group (Yap *et al*., 2020), we decided to focus on the how the restriction of histidine or phenylalanine affects the metabolic health of diet-induced obese mice. We find that restricting histidine dramatically reduces adiposity and restores normal body composition to diet-induced obese mice, while restriction of phenylalanine has a smaller but still positive effect on weight and body composition, and that this is associated with increased energy expenditure. Conversely, we find that dietary histidine levels are correlated with weight and adipose gain during Western diet feeding of young lean mice. Surprisingly, the effects of dietary histidine are independent of the energy balance hormone fibroblast growth factor 21 (FGF21), which has been implicated in the response to protein restriction as well as the restriction of a number of other essential amino acids (Laeger *et al*., 2014; Yap *et al*., 2020; Yu *et al*., 2021). Although restriction of histidine beginning in midlife improves metabolic health in male mice, it does not impact frailty or lifespan in either males or females. Finally, we find that the consumption of dietary protein with increased histidine content is associated with increased body mass index (BMI) in humans. In conclusion, these results demonstrate that dietary histidine is a critical regulator of weight and body composition, and suggest that interventions based on decreasing dietary levels of histidine may be a translatable way to promote and restore metabolic health.

## Results

### Reducing dietary histidine, phenylalanine, or threonine reduces weight and fat mass gain in lean mice

We determined the metabolic effect of restricting each of the six non-BCAA essential amino acids (AAs) individually. Our AA-defined control (Ctrl AA) diet contained all twenty common AAs; the diet composition reflects that of a natural chow diet in which 21% of calories are derived from protein and which we have utilized in several previous studies (Richardson *et al*., 2021; Yu *et al*., 2021). We created a diet series based on this diet, in which either histidine, lysine, methionine, phenylalanine, threonine, or tryptophan was reduced by 67%. All seven diets are isocaloric, with identical levels of fat, and the calories from reduced essential AAs were balanced by increased levels of non-essential AAs to keep the calories derived from AAs constant (**Table 1**).

**Table 1.** Diet composition of diets used to investigate the effect of reduced levels of essential amino acids in the context of a normal calorie diet.

| Amino Acid Diets | Ctrl AA | Low Thr | Low Phe | Low His | Low Met | Low Lys | Low Trp |
| --- | --- | --- | --- | --- | --- | --- | --- |
| Teklad Diet name | 21% Protein | Thr 2/3 Adj Diet | Phe and Tyr Adj Diet | His 2/3 Adj Diet | Met 2/3 Adj Diet | Lys 2/3 Adj Diet | Trp 2/3 Adj Diet |
| Teklad Diet number | TD.140711 | TD.160236 | TD.160235 | TD.160237 | TD.160073 | TD.160075 | TD.160074 |
| Color | Red | Orange | Green | Yellow | Green | Black | Orange |
| Formula | g/kg | g/kg | g/kg | g/kg | g/kg | g/kg | g/kg |
| L-Alanine | 9.38 | 9.9915 | 9.966 | 9.8739 | 9.7175 | 11.0559 | 9.6309 |
| L-Arginine | 6.3 | 6.3 | 6.3 | 6.3 | 6.3 | 6.3 | 6.3 |
| L-Asparagine | 20.58 | 21.0334 | 21.0146 | 20.9462 | 20.8302 | 21.8226 | 20.766 |
| L-Aspartic Acid | 20.58 | 21.4939 | 21.4557 | 21.318 | 21.0843 | 23.0843 | 20.9549 |
| L-Cysteine | 7.2 | 7.2 | 7.2 | 7.2 | 7.2 | 7.2 | 7.2 |
| L-Glutamic Acid | 28.97 | 29.9797 | 29.9377 | 29.7856 | 29.5272 | 31.7374 | 29.3843 |
| L-Glutamine | 33.77 | 34.2759 | 34.2549 | 34.1786 | 34.0492 | 35.1564 | 33.9775 |
| Glycine | 2.96 | 3.475 | 3.4537 | 3.376 | 3.2443 | 4.3718 | 3.1713 |
| L-Histidine HCl, monohydrate | 4.6 | 4.6 | 4.6 | 1.5 | 4.6 | 4.6 | 4.6 |
| L-Isoleucine | 7.8 | 7.8 | 7.8 | 7.8 | 7.8 | 7.8 | 7.8 |
| L-Leucine | 25.4 | 25.4 | 25.4 | 25.4 | 25.4 | 25.4 | 25.4 |
| L-Lysine HCl | 20.38 | 20.38 | 20.38 | 20.38 | 20.38 | 6.64 | 20.38 |
| L-Methionine | 6.7 | 6.7 | 6.7 | 6.7 | 2.18 | 6.7 | 6.7 |
| L-Phenylalanine | 6.6 | 6.6 | 2.15 | 6.6 | 6.6 | 6.6 | 6.6 |
| L-Proline | 7.41 | 8.199 | 8.167 | 8.0479 | 7.8459 | 9.5748 | 7.7341 |
| L-Serine | 7.41 | 8.1311 | 8.1012 | 7.9924 | 7.808 | 9.3864 | 7.7058 |
| L-Threonine | 9.7 | 3.16 | 9.7 | 9.7 | 9.7 | 9.7 | 9.7 |
| L-Tryptophan | 3.4 | 3.4 | 3.4 | 3.4 | 3.4 | 3.4 | 1.1 |
| L-Tyrosine | 6.9 | 6.9 | 2.25 | 6.9 | 6.9 | 6.9 | 6.9 |
| L-Valine | 8.4 | 8.4 | 8.4 | 8.4 | 8.4 | 8.4 | 8.4 |
| Sucrose | 291.248 | 291.248 | 291.248 | 291.248 | 291.248 | 291.248 | 291.248 |
| Corn Starch | 150.0 | 150.5 | 151.9 | 149.3 | 150.7 | 149.3 | 150.0 |
| Maltodextrin | 150.0 | 150.5 | 151.9 | 149.3 | 150.7 | 149.3 | 150.0 |
| Anhydrous Milkfat | 0.0 | 0.0 | 0.0 | 0.0 | 0.0 | 0.0 | 0.0 |
| Cholesterol | 0.0 | 0.0 | 0.0 | 0.0 | 0.0 | 0.0 | 0.0 |
| Corn Oil | 52.0 | 52.0 | 52.0 | 52.0 | 52.0 | 52.0 | 52.0 |
| Olive Oil | 29.0 | 29.0 | 29.0 | 29.0 | 29.0 | 29.0 | 29.0 |
| Cellulose | 30.0 | 30.0 | 30.0 | 30.0 | 30.0 | 30.0 | 30.0 |
| Mineral Mix, AIN-93M-MX (94049) | 35.0 | 35.0 | 35.0 | 35.0 | 35.0 | 35.0 | 35.0 |
| Calcium Phosphate Ca(H<sub>2</sub>PO<sub>4</sub>)<sub>2</sub> · H<sub>2</sub>O | 8.2 | 8.2 | 8.2 | 8.2 | 8.2 | 8.2 | 8.2 |
| Vitamin Mix, Teklad (40060) | 10.0 | 10.0 | 10.0 | 10.0 | 10.0 | 10.0 | 10.0 |
| TBHQ, antioxidant | 0.012 | 0.012 | 0.012 | 0.012 | 0.012 | 0.012 | 0.012 |
| Food Coloring | 0.1 | 0.1 | 0.1 | 0.1 | 0.1 | 0.1 | 0.1 |
| % kcal from |  |  |  |  |  |  |  |
| Protein (based on N x 6.25) | 22 | 22 | 22 | 24.1 | 23.8 | 24.2 | 23.9 |
| Carbohydrates | 59.4 | 59.4 | 58.3 | 57.8 | 58.2 | 57.6 | 57.9 |
| Fat | 18.6 | 18.6 | 18.2 | 18.2 | 18.2 | 18.1 | 18.2 |
| Kcal/g | 3.9 | 3.9 | 4 | 4 | 4 | 4 | 3.9 |

In our first experimental cohort, we fed C57BL/6J male mice either a Ctrl AA diet or a diet in which either threonine (Low Thr), phenylalanine (Low Phe), or histidine (Low His) was reduced by 67% (**Fig. 1A**). Tracking weight and body composition over three months, we observed significant effects of Low Phe and Low His diets, with mice fed these diets gaining less than half the weight of Ctrl AA-fed mice (**Fig. 1B**). The effect of the Low Phe and Low His diets was due to a reduction in both fat mass and lean mass gain (**Figs. 1C-1D**). In contrast, a Low Thr diet had no significant effect on weight, but had significantly reduced fat mass gain with a numerical increase in lean mass. The overall effect of all three diets was decreased adiposity (**Fig. 1E**).

**Figure 1.**
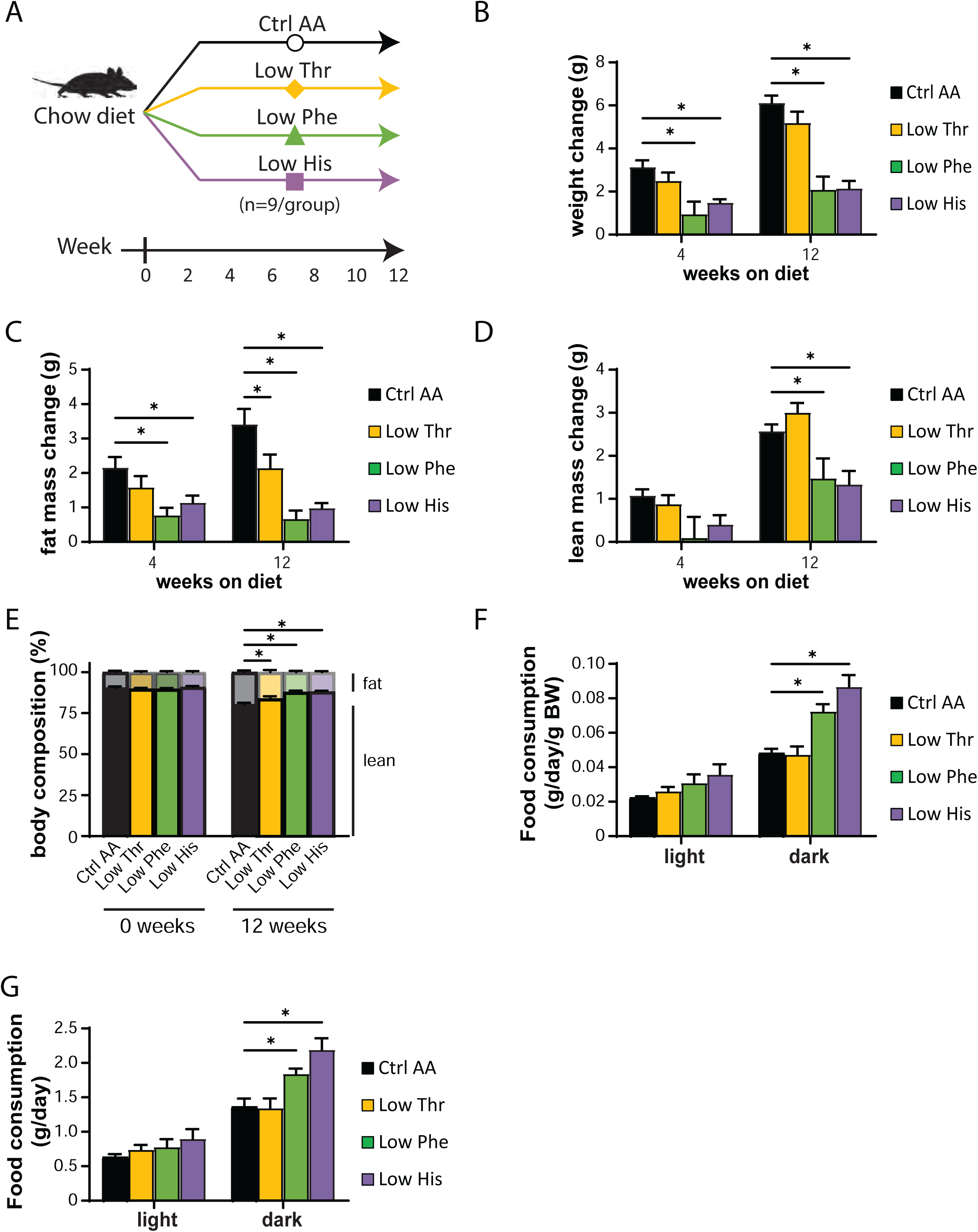
Dietary restriction of histidine, phenylalanine or threonine improves body composition. (A) Experimental scheme. (B-E) Change in body weight (B), fat mass (C), lean mass (D), and body composition (E) of mice consuming the indicated diets for the indicated time (n=8-9 mice per group, *p<0.05, Dunnett’s test vs. Ctrl AA following a mixed-effects model (Restricted Maximum Likelihood (REML)). (F-G) Food consumption (F) and energy expenditure (heat) (G) normalized to body weight of mice fed the indicated diets (n=8-9 per group, *p<0.05, Dunnet’s test vs. Ctrl AA conducted separately for the light and dark cycles post 2-way RM ANOVA). Data represented as mean ± SEM.

We used metabolic chambers to evaluate energy balance, analyzing food consumption, spontaneous activity, and energy expenditure. Intriguingly Low Phe-fed and Low His-fed mice had a significant increase in food consumption on both a per mouse and per body weight basis (**Figs. 1F-1G**). Consistent with the reduced weight gain and fat accretion of these mice, this increased calorie intake was offset by increased energy expenditure, while spontaneous activity was unaltered (**Figs. 2A-2C**). The respiratory exchange ratio (RER), which reflects substrate utilization, was increased at night in mice consuming Low Phe or Low His diets, suggesting increased carbohydrate utilization (**Fig. 2D**). Finally, we assessed blood glucose control by conducting glucose and insulin tolerance tests. We observed no significant effects of any of the diets on either glucose tolerance or insulin sensitivity (**Figs. 2E-2F**).

**Figure 2.**
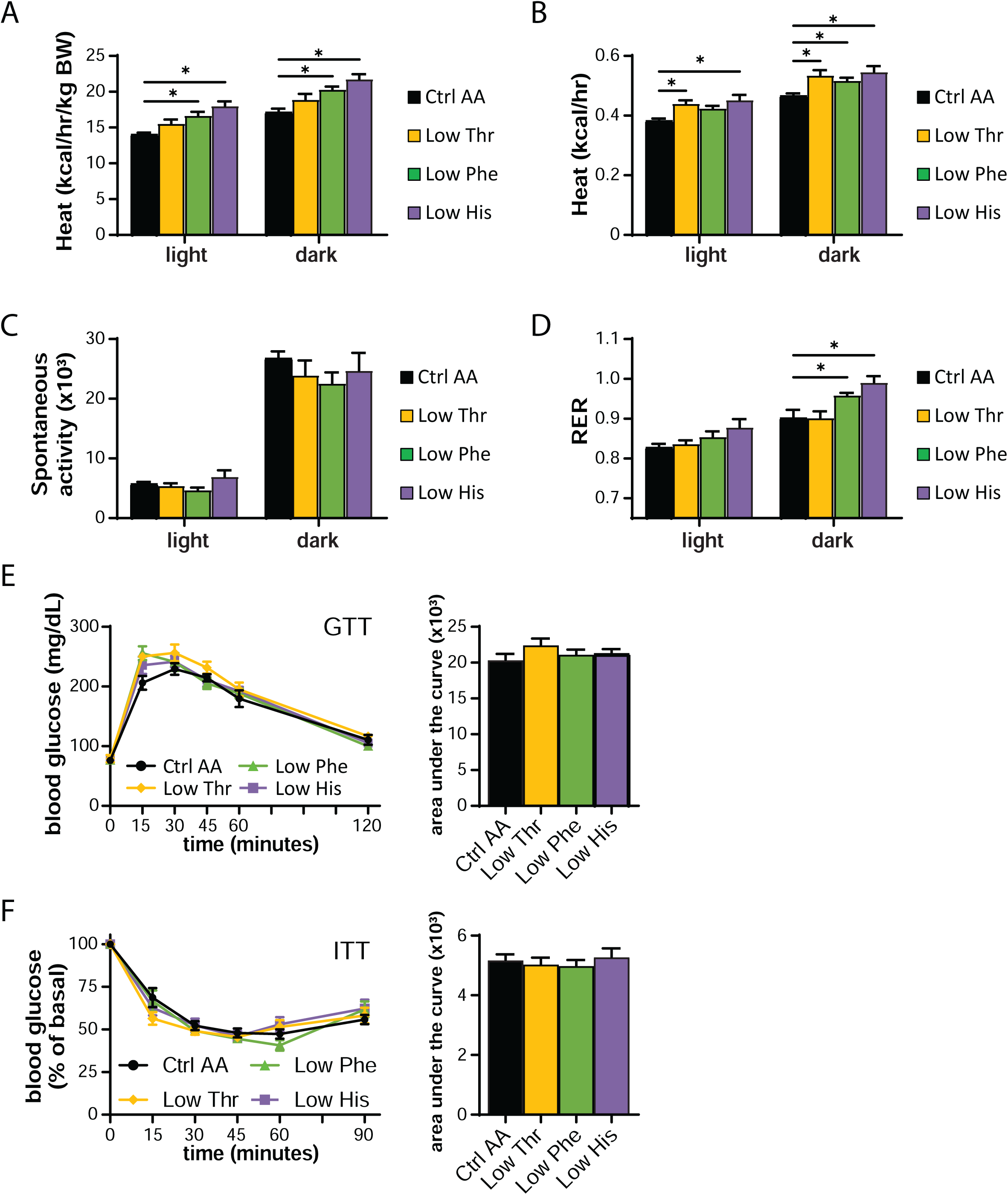
Dietary restriction of histidine, phenylalanine or threonine alters energy balance but not glycemic control. (A-D) Metabolic chambers were utilized to assess multiple components of energy balance, including energy expenditure (heat) (A-B), spontaneous activity (C), and respiratory exchange ratio (RER) (D); n=3-6 per group, *p<0.05, Dunnet’s test vs. WD Ctrl AA conducted separately for the light and dark cycles post 2-way RM ANOVA. (E) Glucose tolerance test (n=9/group, *p<0.05 vs. Ctrl AA, Dunnett’s test post ANOVA). (F) Insulin tolerance test (n=8-9/group, *p<0.05 vs. Ctrl AA, Dunnett’s test post ANOVA). Data represented as mean ± SEM.

### Reducing lysine, methionine, or tryptophan has minimal effects on weight and body composition

We utilized a second cohort of mice to test the effect of specifically reducing methionine (Low Met), lysine (Low Lys), or tryptophan (Low Trp) by 67% (**Fig. 3A**). We found that these diets had minimal effects on weight, fat and lean mass accretion, or adiposity, with weight and no other parameters being affected by a Low Trp diet only (**Figs. 3B-3E**). Consistent with these findings, food consumption, energy expenditure, and substrate utilization was not altered in mice fed any of these three diets (**Figs. 3F-3G and 4A-4D**). We again observed no significant effects of any of the diets on either glucose tolerance or insulin sensitivity (**Figs. 4E-4F**).

**Figure 3.**
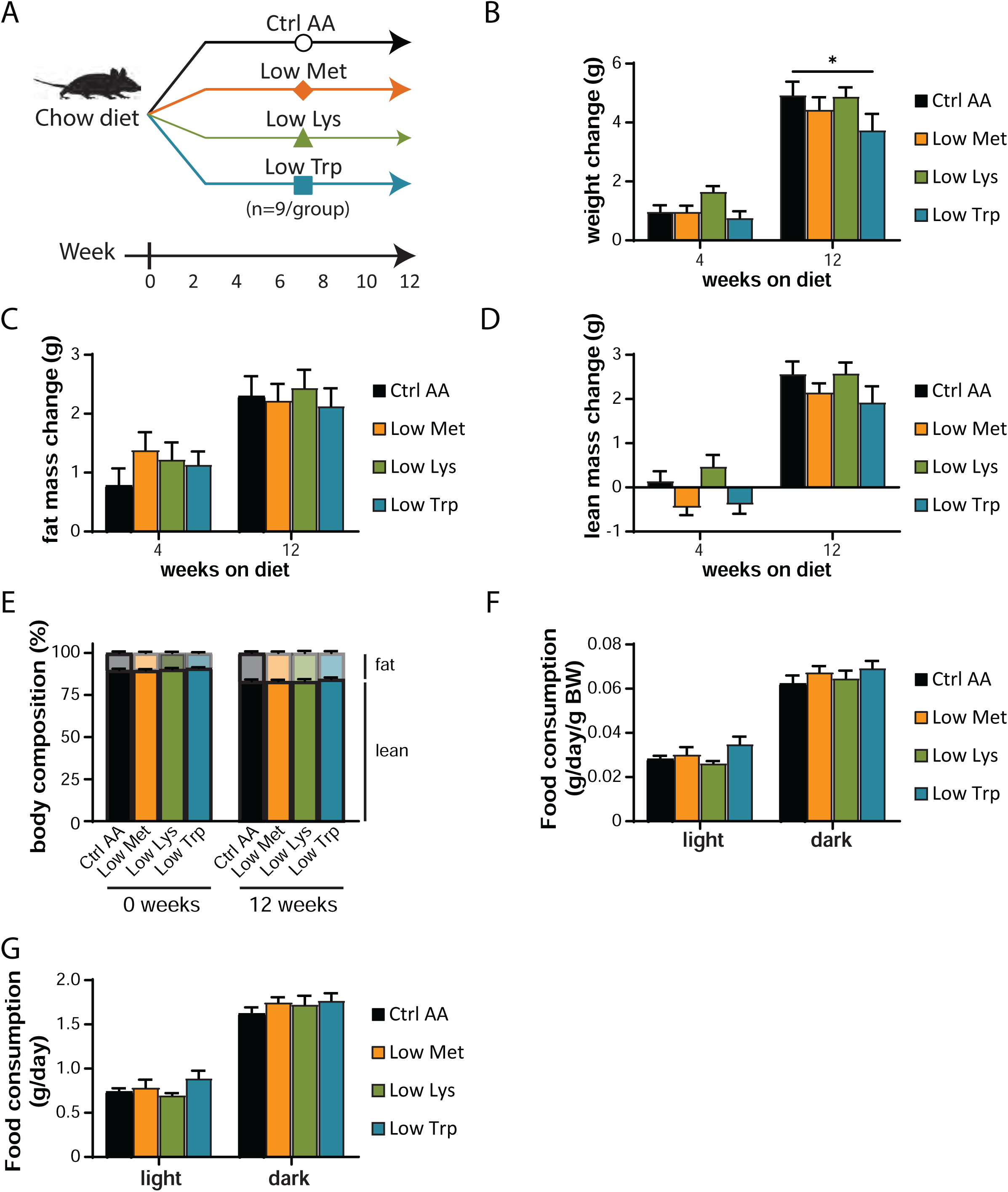
Dietary restriction of lysine, methionine, or tryptophan has no effect on body composition. (A) Experimental scheme. (B-E) Change in body weight (B), fat mass (C), lean mass (D), and body composition (E) of mice consuming the indicated diets for the indicated time (n=9 mice per group, *p<0.05, Dunnett’s test vs. Ctrl AA post 2-way RM ANOVA). (F-G) Food consumption (F) and energy expenditure (heat) (G) normalized to body weight of mice fed the indicated diets (n=8-9 per group, *p<0.05, Dunnet’s test vs. Ctrl AA conducted separately for the light and dark cycles post 2-way RM ANOVA). Data represented as mean ± SEM.

**Figure 4.**
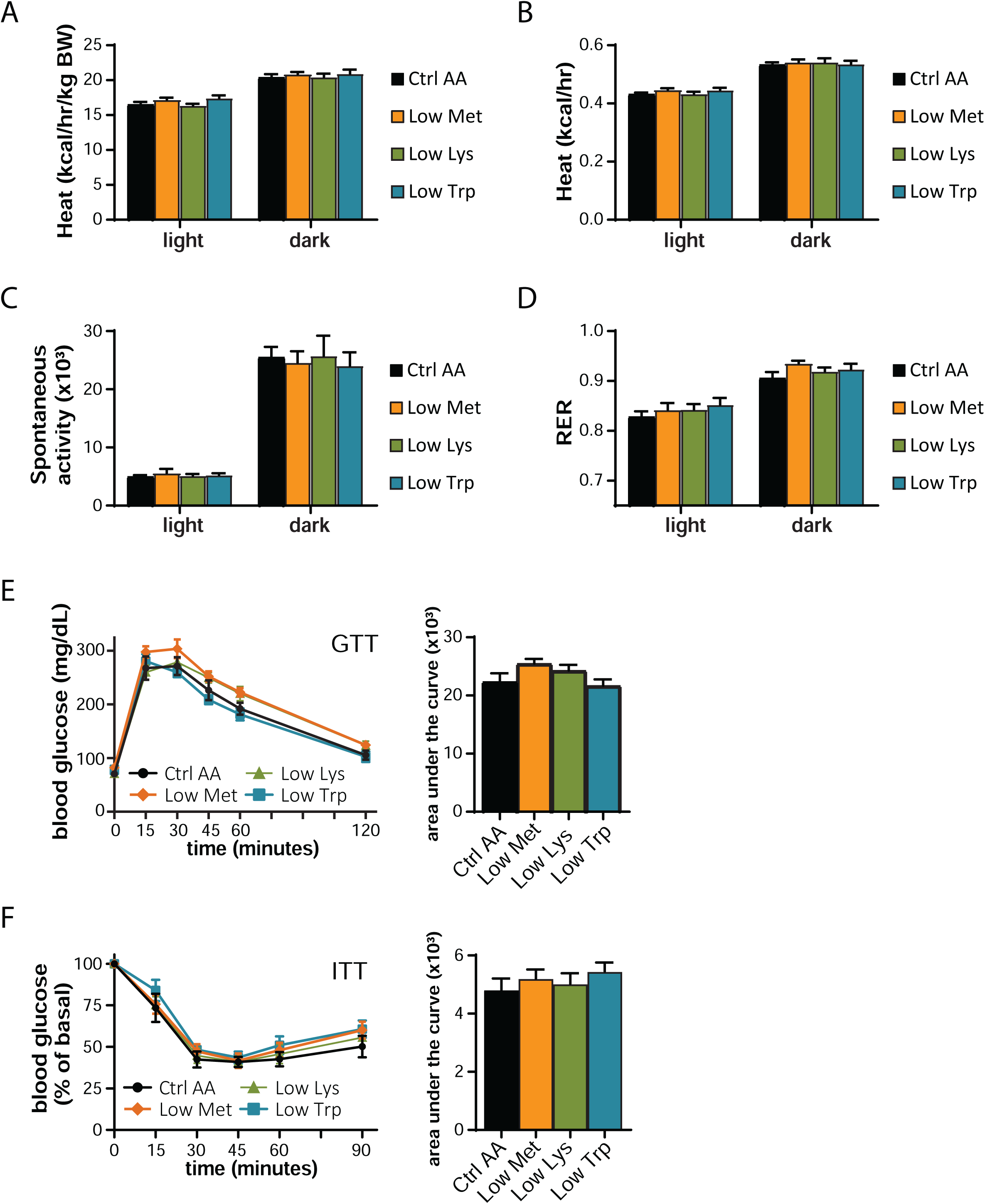
Dietary restriction of lysine, methionine, or tryptophan has no effect on energy balance or glycemic control. (A-D) Metabolic chambers were utilized to assess multiple components of energy balance, including, energy expenditure (heat) (A-B), spontaneous activity (C), and respiratory exchange ratio (RER) (D); n=8-9 per group, *p<0.05, Dunnet’s test vs. WD Ctrl AA conducted separately for the light and dark cycles post 2-way RM ANOVA. (E) Glucose tolerance test (n=9/group, *p<0.05 vs. Ctrl AA, Dunnett’s test post ANOVA). (F) Insulin tolerance test (n=9/group, *p<0.05 vs. Ctrl AA, Dunnett’s test post ANOVA). Data represented as mean ± SEM.

### Specifically restricting dietary histidine restores normal body composition to DIO mice, increasing energy expenditure

We chose to focus our efforts on phenylalanine and histidine, as restriction of these two essential AAs had by far the greatest impact on weight, body composition, and energy balance; additionally, another group had already found beneficial metabolic effects of restricting dietary threonine (Yap *et al*., 2020). In our previous work, we determined that restriction of dietary protein can restore metabolic health to diet-induced obese (DIO) even while they continue to consume a high-fat, high-sucrose Western Diet (WD) (Cummings *et al*., 2018; Yu *et al*., 2021). We therefore decided to examine the metabolic impact of reducing levels of phenylalanine or histidine in mice pre-conditioned with a WD for 12 weeks (**Fig. 5A**).

**Figure 5.**
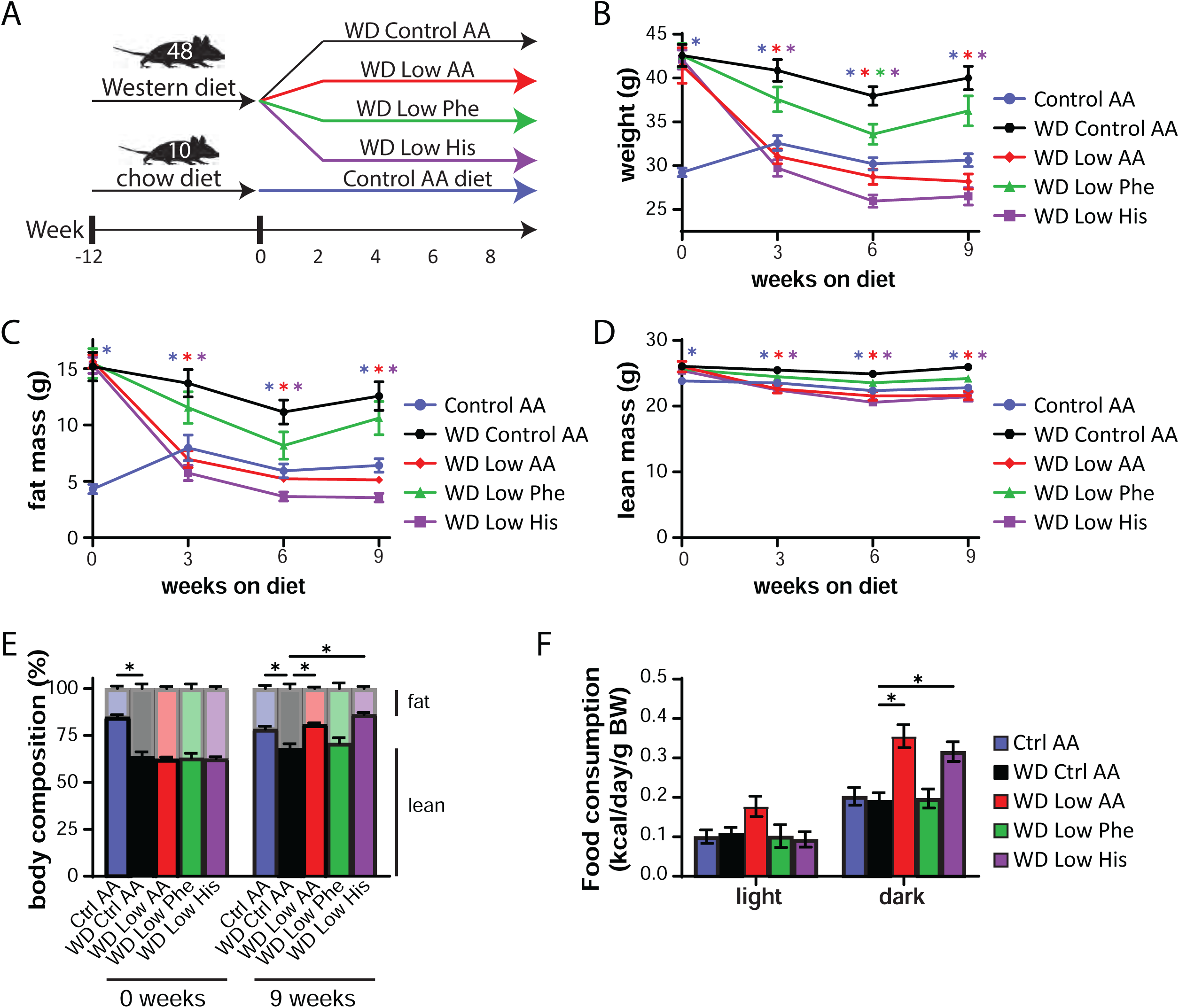
Restricting dietary histidine improves the metabolic health of DIO mice. (A) Experimental scheme. (B-E) Body weight (B), fat mass (C), lean mass (D), and body composition (E) of mice consuming the indicated diets for the indicated time (n=10-12 mice per group, *p<0.05, Dunnett’s test vs. WD Ctrl AA post 2-way RM ANOVA). (F-G) Metabolic chambers were used to assess multiple components of energy balance including food consumption normalized to body weight (n=5-8 per group, *p<0.05, Dunnet’s test vs. WD Ctrl AA conducted separately for the light and dark cycles post 2-way RM ANOVA).

To specifically address this question, we designed a new series of diets based on an amino acid-defined WD (WD Control AA) that we have previously utilized (Cummings *et al*., 2018; Richardson *et al*., 2021); this diet closely matches the macronutrient profile of a widely used naturally sourced Western diet (TD.88137). Briefly, the level of either phenylalanine (WD Low Phe), histidine (WD Low His), or all AAs (WD Low AA) was reduced by 67% in the context of a WD; all of the WDs were isocaloric with identical levels of fat, and in case of the WD Low Phe and WD Low His diets, the percentage of calories derived from AAs was kept constant by proportionally adjusting the amount of non-essential AAs (**Table 2**). In parallel, a group of chow-fed mice never exposed to a WD were switched to the Ctrl AA diet.

**Table 2.** Diet composition of diets used to investigate the effect of reduced levels of essential amino acids in the context of a WD.

| Amino Acid Defined Diets | Ctrl AA | WD Ctrl AA | WD Low AA | WD Low Phe | WD Low His |
| --- | --- | --- | --- | --- | --- |
| Teklad Diet name | 21% Protein | AA Adj<br>Calories, 21%<br>Milkfat, 0.2%<br>Chol | AA Adj<br>Calories, 7%<br>Protein, 21%<br>Milkfat | Phe and Tyr<br>Adj Western<br>(Milkfat, Chol,<br>Red) | His 2/3 Adj<br>Western<br>(Milkfat, Chol,<br>Green) |
| Teklad Diet number | TD.140711 | TD.160186 | TD.160187 | TD.171009 | TD.171010 |
| Color | Red | Aqua | Orange | Red | Green |
| Formula | g/kg | g/kg | g/kg | g/kg | g/kg |
| L-Alanine | 9.38 | 9.38 | 3.05 | 9.966 | 9.8739 |
| L-Arginine | 6.3 | 6.3 | 2.05 | 6.3 | 6.3 |
| L-Asparagine | 20.58 | 20.58 | 6.7 | 21.1046 | 20.9462 |
| L-Aspartic Acid | 20.58 | 20.58 | 6.7 | 21.4557 | 21.318 |
| L-Cysteine | 7.2 | 7.2 | 2.34 | 7.2 | 7.2 |
| L-Glutamic Acid | 28.97 | 28.97 | 9.43 | 29.9377 | 29.79 |
| L-Glutamine | 33.77 | 33.77 | 11 | 34.2549 | 34.18 |
| Glycine | 2.96 | 2.96 | 0.96 | 3.4537 | 3.38 |
| L-Histidine HCl, monohydrate | 4.6 | 4.6 | 1.5 | 4.6 | 1.5 |
| L-Isoleucine | 7.8 | 7.8 | 2.54 | 7.8 | 7.8 |
| L-Leucine | 25.4 | 25.4 | 8.27 | 25.4 | 25.4 |
| L-Lysine HCl | 20.38 | 20.38 | 6.64 | 20.38 | 20.38 |
| L-Methionine | 6.7 | 6.7 | 2.18 | 6.7 | 6.7 |
| L-Phenylalanine | 6.6 | 6.6 | 2.15 | 2.15 | 6.6 |
| L-Proline | 7.41 | 7.41 | 2.41 | 8.167 | 8.05 |
| L-Serine | 7.41 | 7.41 | 2.41 | 8.1012 | 7.99 |
| L-Threonine | 9.7 | 9.7 | 3.16 | 9.7 | 9.7 |
| L-Tryptophan | 3.4 | 3.4 | 1.1 | 3.4 | 3.4 |
| L-Tyrosine | 6.9 | 6.9 | 2.25 | 2.25 | 6.9 |
| L-Valine | 8.4 | 8.4 | 2.735 | 8.4 | 8.4 |
| Sucrose | 291.248 | 341.46 | 341.46 | 341.46 | 341.46 |
| Corn Starch | 150.0 | 49.63 | 132.0625 | 51.5346 | 48.9509 |
| Maltodextrin | 150.0 | 49.63 | 132.0625 | 51.5346 | 48.9509 |
| Anhydrous Milkfat | 0.0 | 210 | 210 | 210 | 210 |
| Cholesterol | 0.0 | 1.5 | 1.5 | 1.5 | 1.5 |
| Corn Oil | 52.0 | 0.0 | 0.0 | 0.0 | 0.0 |
| Olive Oil | 29.0 | 0.0 | 0.0 | 0.0 | 0.0 |
| Cellulose | 30.0 | 50 | 50 | 50 | 50 |
| Mineral Mix, AIN-93M-MX (94049) | 35.0 | 35 | 35 | 35 | 35 |
| Calcium Phosphate Ca(H<sub>2</sub>PO<sub>4</sub>)<sub>2</sub> · H<sub>2</sub>O | 8.2 | 8.2 | 8.2 | 8.2 | 8.2 |
| Vitamin Mix, Teklad (40060) | 10.0 | 10 | 10 | 10 | 10 |
| TBHQ, antioxidant | 0.012 | 0.04 | 0.04 | 0.04 | 0.04 |
| Food Coloring | 0.1 | 0.1 | 0.1 | 0.1 | 0.1 |
| % kcal from |  |  |  |  |  |
| Protein (based on N x 6.25) | 22 | 20.7 | 6.8 | 18.9 | 19 |
| Carbohydrates | 59.4 | 38.5 | 52 | 39.5 | 39.2 |
| Fat | 18.6 | 40.9 | 41.2 | 41.6 | 41.8 |
| Kcal/g | 3.9 | 4.6 | 4.6 | 4.5 | 4.5 |

As we expected based on our previous studies, DIO mice switched to the WD Control AA diet remained obese, lean mice switched to the Ctrl AA diet remained comparatively light and lean, and mice switched to the WD Low AA diet rapidly lost weight and fat mass, with an overall reduction in adiposity (**Fig. 5B-5E**). Surprisingly, DIO mice switched to the WD Low His diet diets progressively lost weight for 3 weeks, similarly to the WD Low AA-fed mice; both the WD Low His and WD Low AA-fed mice then continued to lose weight at a reduced weight for the next 3 weeks, which then stabilized (**Fig. 5B**). The majority of the weight loss of both groups of mice was due to a dramatic decrease in fat mass, whereas lean mass was not as affected, leading to an overall reduction in adiposity (**Fig. 5C-5E**). WD Low Phe-fed mice showed a lesser degree of weight loss, and did not have a statistically significant different in fat mass or adiposity from mice fed the WD Control AA diet (**Fig. 5B-5E**).

We next used metabolic chambers to evaluate energy balance, analyzing food consumption, spontaneous activity, and energy expenditure. We observed that mice fed either the WD Low AA and WD Low His diet had higher food consumption than WD Control AA-fed mice on both a per mouse and per body weight basis (**Figs. 5F-5G**). Consistent with the reduced body weight of animals fed these diets, our previous observations of WD Low AA-fed mice (Cummings *et al*., 2018; Yu *et al*., 2021), and our observations in lean Low His-fed mice not exposed to a WD, we found that the WD Low AA-fed and WD Low His-fed mice had increased energy expenditure normalized to body weight (**Figs. 6A-6B)**. This occurred despite a decrease in activity during the dark cycle that was statistically significant for the WD Low AA-fed mice (**Fig. 6C**). Similar to our findings in lean mice and previous observations of WD Low AA-fed mice, we observed that RER was increased during the dark cycle in mice consuming the WD Low AA and WD Low His diets (**Fig. 6D**). Not surprisingly given the minor beneficial impact of a WD Low Phe diet on weight and body composition, the only effect of a WD Low Phe diet on components of energy balance that reached statistical significance was a small increase in energy expenditure normalized to body weight (**Fig. 6A**).

**Figure 6.**
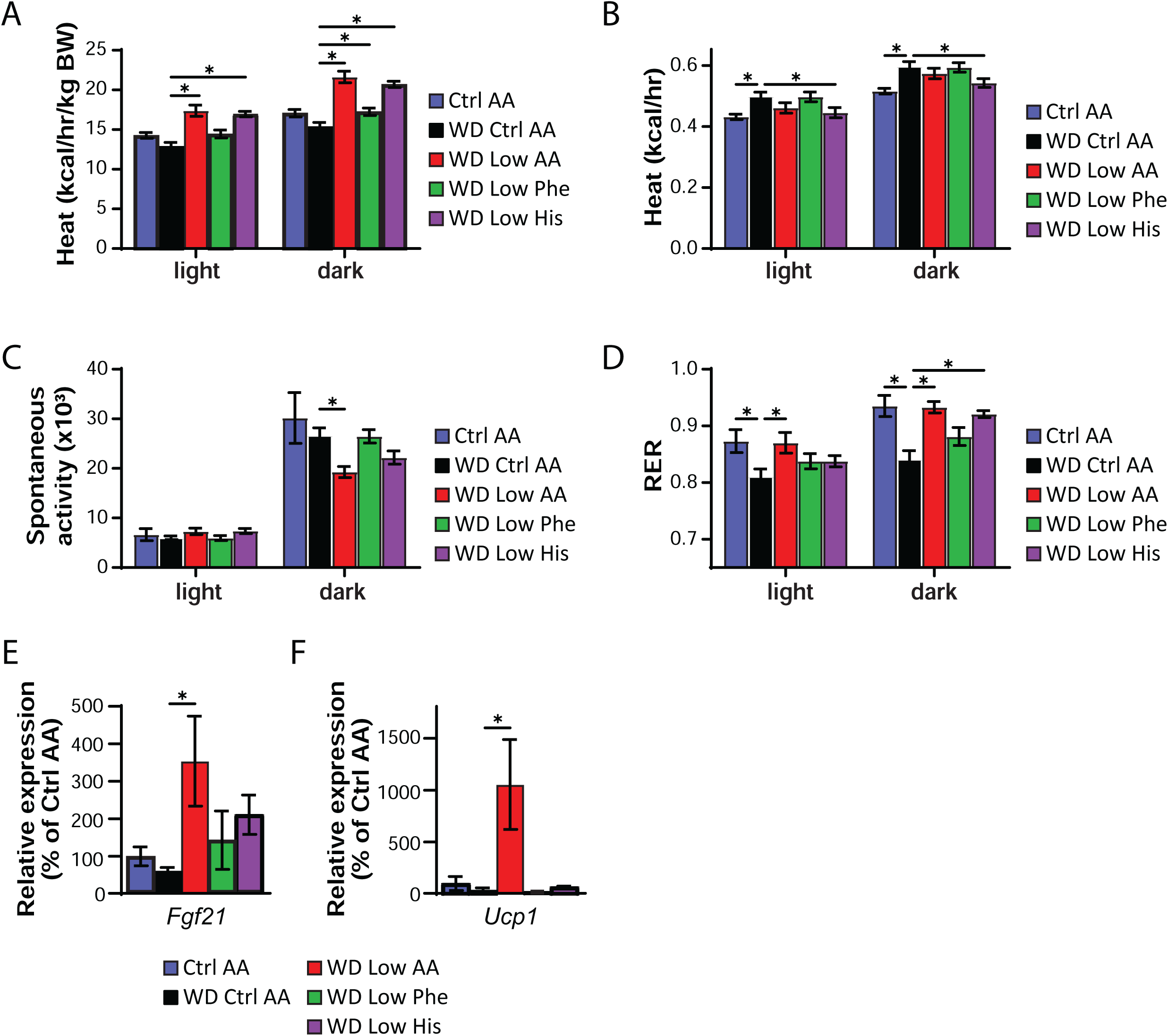
Restricting dietary histidine alters energy balance of diet-induced obese mice. (A-D) Metabolic chambers were utilized to assess multiple components of energy balance, including energy expenditure (heat) (A-B), spontaneous activity (C), and respiratory exchange ratio (RER) (D); n=8/group, *p<0.05, Dunnet’s test vs. WD Ctrl AA conducted separately for the light and dark cycles post 2-way RM ANOVA. (E-F) *Fgf21* expression in the liver (E) and *Ucp1* expression in the iWAT (F) of mice fed the indicated diets for 9 weeks (n = 6-9 per group, *p<0.05, Dunnett’s test vs. WD Ctrl AA post ANOVA). Data represented as mean ± SEM.

Protein restriction induces hepatic expression of the energy balance hormone *Fgf21* (Laeger *et al*., 2014), and many of the metabolic benefits of PR, including increased energy, in C57BL/6J male mice are mediated by FGF21 through induction of uncoupling protein 1 (UCP1) in inguinal white adipose tissue (iWAT) (Hill *et al*., 2017; Green *et al*., 2022). In agreement with our observation of increased energy expenditure in WD Low AA-fed mice, we observed increased hepatic expression of *Fgf21* and iWAT expression of *Ucp1* in these animals (**Figs. 6E-6F**). However, neither hepatic *Fgf21* nor iWAT *Ucp1* was elevated in WD Low His-fed mice.

### Restricting dietary histidine improves glycemic control and hepatic health of DIO mice

We assessed blood glucose control by conducting intraperitoneal glucose and insulin tolerance tests. We observed that mice consuming the WD Low AA and WD Low His diets had improved glucose tolerance relative not only to WD Control AA-fed mice, but to Ctrl AA-fed mice of a similar weight never exposed to a WD (**Fig. 7A**). Mice consuming the WD Low AA, WD Low His, and WD Low Phe diets all had improved insulin tolerance relative to WD Control AA- fed mice (**Fig. 7B**).

**Figure 7.**
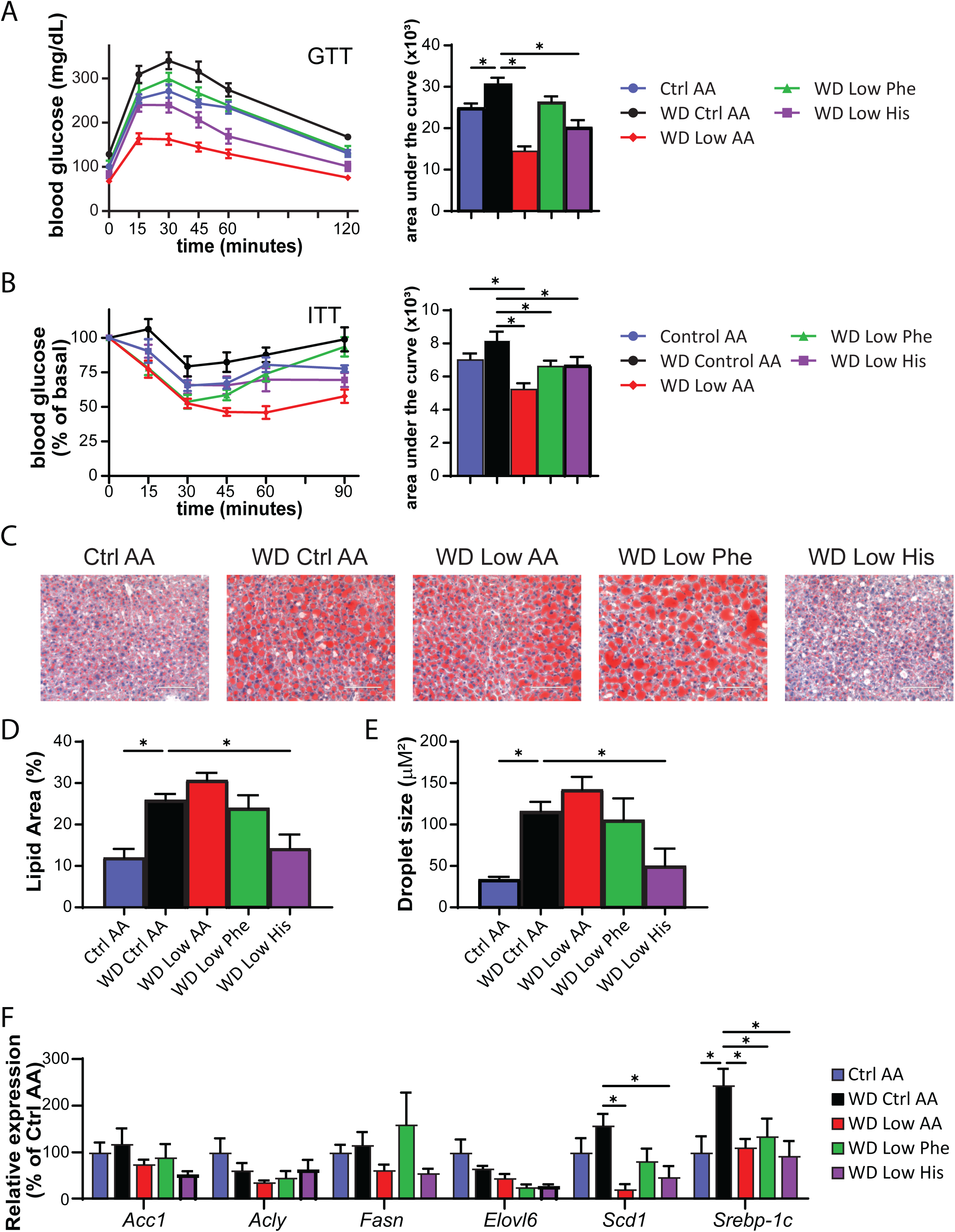
Restricting dietary histidine improves glycemic control and hepatic steatosis of DIO mice. (A) Glucose tolerance test after 3 weeks on the indicated diets (n=10-12/group, *p<0.05 vs. WD Ctrl AA, Dunnett’s test post ANOVA). (B) Insulin tolerance test after 4 weeks on the indicated diets (n=9-12/group, *p<0.05 vs. WD Ctrl AA, Dunnett’s test post ANOVA). (C-E) Representative Oil-Red-O-stained sections of liver of mice after 9 weeks on the indicated diets. Scale bar, 100 µm (C); quantification of lipid droplet area (D) and lipid droplet size (E) (n=6/group, *p<0.05 vs. WD Ctrl AA, Dunnett’s test post ANOVA). (F) Lipogenic gene expression after 9 weeks on the indicated diets (n=6/group, *p<0.05 vs. WD Ctrl AA, Dunnett’s test post 2-way RM ANOVA). Data represented as mean ± SEM.

WD feeding induces hepatic steatosis, likely contributing to the negative effects of a WD on glucose tolerance. As expected, WD Ctrl AA fed mice had evident hepatic steatosis and large lipid droplets, whereas the mice switched to the WD Low His diet had decreased liver droplet size and normal liver histology at the conclusion of the experiment (**Figs. 7C-7E**). As we have previously observed, restriction of all AAs (WD Low AA diet) did not lead to improved histology or smaller lipid droplets. We quantified hepatic expression of several lipogenic genes as well as the transcription factor *Srebp-1c*. We observed numerical decreases in the mRNA expression of four lipogenic genes, *Acc1*, *Fasn*, *Elovl6*, and *Scd1* in WD Low His-fed mice relative to WD Ctrl AA-fed mice; the decrease was statistically significant in the case of *Scd1*, although this did not likely account for the specific effect of the WD Low His diet since it also occurred with total AA restriction (**Fig. 7F**). Similarly, expression of the transcription factor *Srebp-1c* was also significantly reduced in WD Low His-fed and WD Low AA-fed mice (**Fig. 7F**).

To quantitatively assess how dietary histidine levels impacted hepatic insulin sensitivity and glucose uptake by peripheral tissues, we performed a hyperinsulinemia-euglycemic clamp on DIO mice switched to either a WD Ctrl AA or WD Low His diet (**Fig. 8A–8J**). Mice fed the WD Low His diet required a considerably greater glucose infusion rate (GIR) to achieve euglycemia than mice fed the WD Ctrl AA (**Fig. 8A**). Calculating hepatic glucose production under basal and hyperinsulinemic conditions revealed that there was no difference in hepatic insulin sensitivity between WD Low His-fed mice and WD Ctrl AA-fed mice (**Figs. 8B-8C**). Thus, the greater GIR required for WD Low His-fed mice to achieve euglycemia was entirely the effect of increased peripheral glucose disposal under clamp conditions (**Fig. 8D**). WD Low His-fed mice had significantly increased glucose uptake rates into critical metabolic tissues, including iWAT, brown adipose tissue (BAT), heart, and skeletal muscle (**Figs. 8E-8H**). In contrast, there were no significant differences in glucose uptake into epididymal white adipose tissue (eWAT) or brain (**Figs. 8I-8J**).

**Figure 8.**
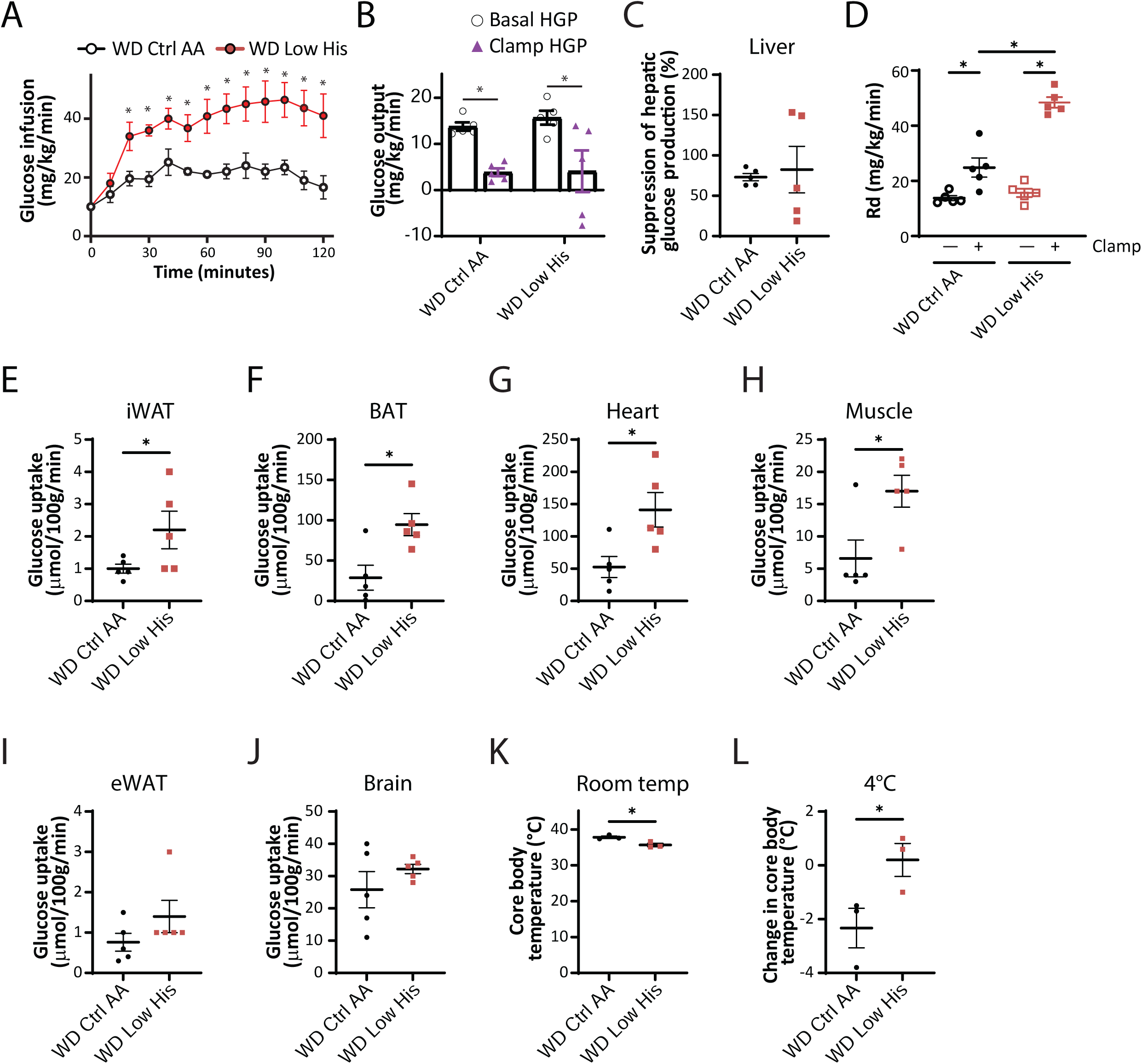
Dietary restriction of histidine increases glucose uptake into multiple tissues of DIO mice. (A-C) Glucose infusion rate (A), basal and clamp hepatic glucose production (B), insulin responsiveness (C) and glucose disposal rate (D) were determined during a hyperinsulinemic-euglycemic clamp in mice preconditioned with a WD for 12 weeks and then switched to a WD Ctrl AA diet or a WD Low His diet for 4 weeks (n = 5/group; (A,C) *p<0.05, t-test; (B, D) *p < 0.05, Sidak’s test post 2-way ANOVA). (E-J) Glucose uptake into iWAT (E), BAT (F), heart (G), muscle (H), eWAT (I), and brain (J) were determined during a hyperinsulinemic-euglycemic clamp in mice preconditioned with a WD for 12 weeks and then switched to a WD Ctrl AA diet or a WD Low His diet for 4 weeks (n = 5/group; *p<0.05, t-test). (K-L) Core temperature was determined using a rectal thermometer at room temp (24°C) (K) and after 1 hour at 4°C (L); n = 3/group, *p < 0.05, t-test. WD Ctrl AA clamp and glucose uptake data has been previously published (Yu *et al*., 2021). Data represented as mean ± SEM.

We considered the possibility that the greater energy expenditure of WD Low His-fed mice and the greater glucose uptake into iWAT and BAT of these animals were linked to increased thermogenesis. We observed a small, statistically significant difference in body temperature, with WD Low His-fed mice having a lower core body temperature than WD Ctrl AA-fed mice (**Fig. 8K**). However, WD Low His-fed mice showed resistance to cold stress, maintaining a significantly higher core body temperature upon exposure to 4°C **(Fig. 8L).**

### Benefits of histidine restriction do not require FGF21

As noted above many of the metabolic benefits of PR, including increased energy, in C57BL/6J male mice are mediated by the hormone FGF21, including increased food consumption and energy expenditure (Hill *et al*., 2017; Green *et al*., 2022). Although we did not observe a statistically significance increase in liver *Fgf21* in histidine restricted animals, we decided to directly test the role of FGF21 in the response to histidine restriction by comparing the response of wild-type and *Fgf21^-/-^* mice to a Low His diet. We placed wild-type (WT) and *Fgf21^-/-^* (KO) mice on either Ctrl AA or Low His diets, and then analyzed weight and body composition. We found that deletion of *Fgf21* did not block the effects of a Low His diet on body weight, fat mass, or lean mass (**Figs. 9A-9C**).

**Figure 9.**
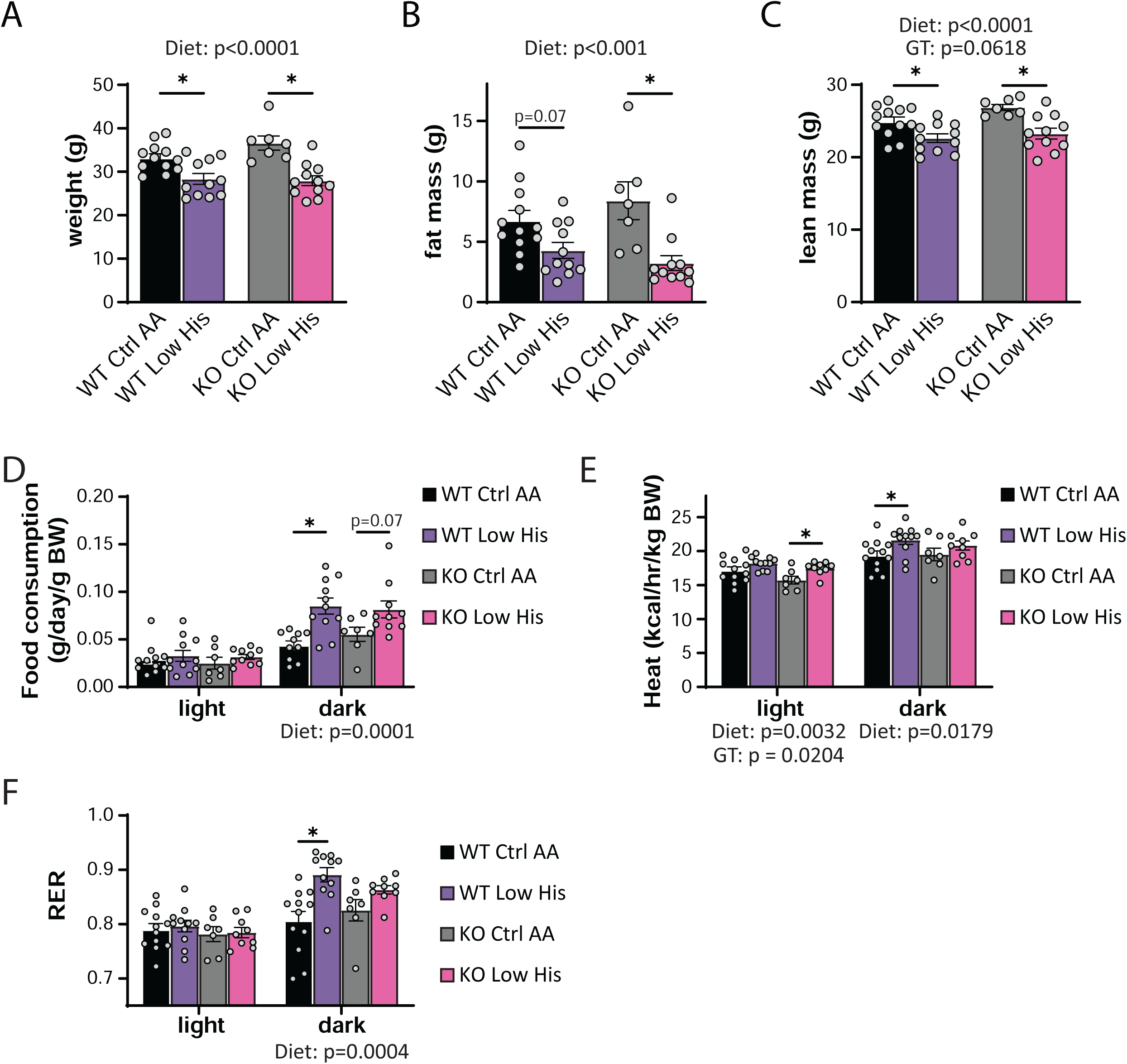
FGF21 is dispensable for the metabolic response to histidine restriction. (A-C) Weight (A), fat mass (B), and lean mass (C), and fat mass of wild-type WT and *Fgf21^-/-^* (KO) mice on the indicated diets (n=7-12 per group, the overall effect of genotype [GT], diet, and the interaction represent the p value from a two-way ANOVA; *p<0.05, Sidak’s test post two-way ANOVA). (D-F) Food consumption normalized to body weight (D), energy expenditure normalized to body weight (E), and RER (F) of mice fed the indicated diets (n=7-11 per group, the overall effect of genotype [GT], diet, and the interaction represent the p value from a two-way ANOVA conducted separately for the light and dark cycles; *p < 0.05, Sidak’s test post two-way ANOVA). Data represented as mean ± SEM.

We next assessed food intake, energy expenditure, and fuel source utilization using metabolic chambers. As expected, we observed a significant overall effect of diet on food consumption, with WT and KO mice responding similarly and increasing food consumption during the dark cycle when fed a Low His diet (**Fig. 9D**). Energy expenditure likewise showed a significant effect of diet in both the light and dark phases, with both WT and KO mice increasing energy expenditure when fed a Low His diet; there was also an overall main effect of genotype on energy expenditure, with KO mice expending slightly less energy during the light phase (**Fig. 9E**). Fuel source utilization, as reflected by changes in the respiratory exchange ratio, was also altered by a Low His diet during the dark cycle in both WT and KO mice (**Fig. 9F**). While there was not a significant interaction between diet and genotype with respect to food consumption, energy expenditure, or RER, in all three cases the response of KO mice was blunted compared to the response seen in WT mice.

### Restriction of histidine starting in midlife improves the metabolic health of males but does not increase lifespan

Restriction of dietary protein and specific AAs, including methionine and the branched-chain AAs, has been shown to promote healthspan and longevity in rodents. To examine the effects of reducing dietary histidine starting in midlife, we randomized 16-month-old male and female C57BL/6.Nia mice from the National Institute on Aging (NIA) Aged Mouse Colony to the Ctrl AA or Low His diet. We followed these animals longitudinally, with assessment of metabolic health and frailty as they aged, and determined their survival. While the body composition of aged female mice was not significantly altered by consumption of the Low His diet, we found that there was a significant effect of a Low His diet on the fat mass and lean mass of male mice; the overall effect was one of reduced male adiposity (**Figs. 10A-D**).

**Figure 10.**
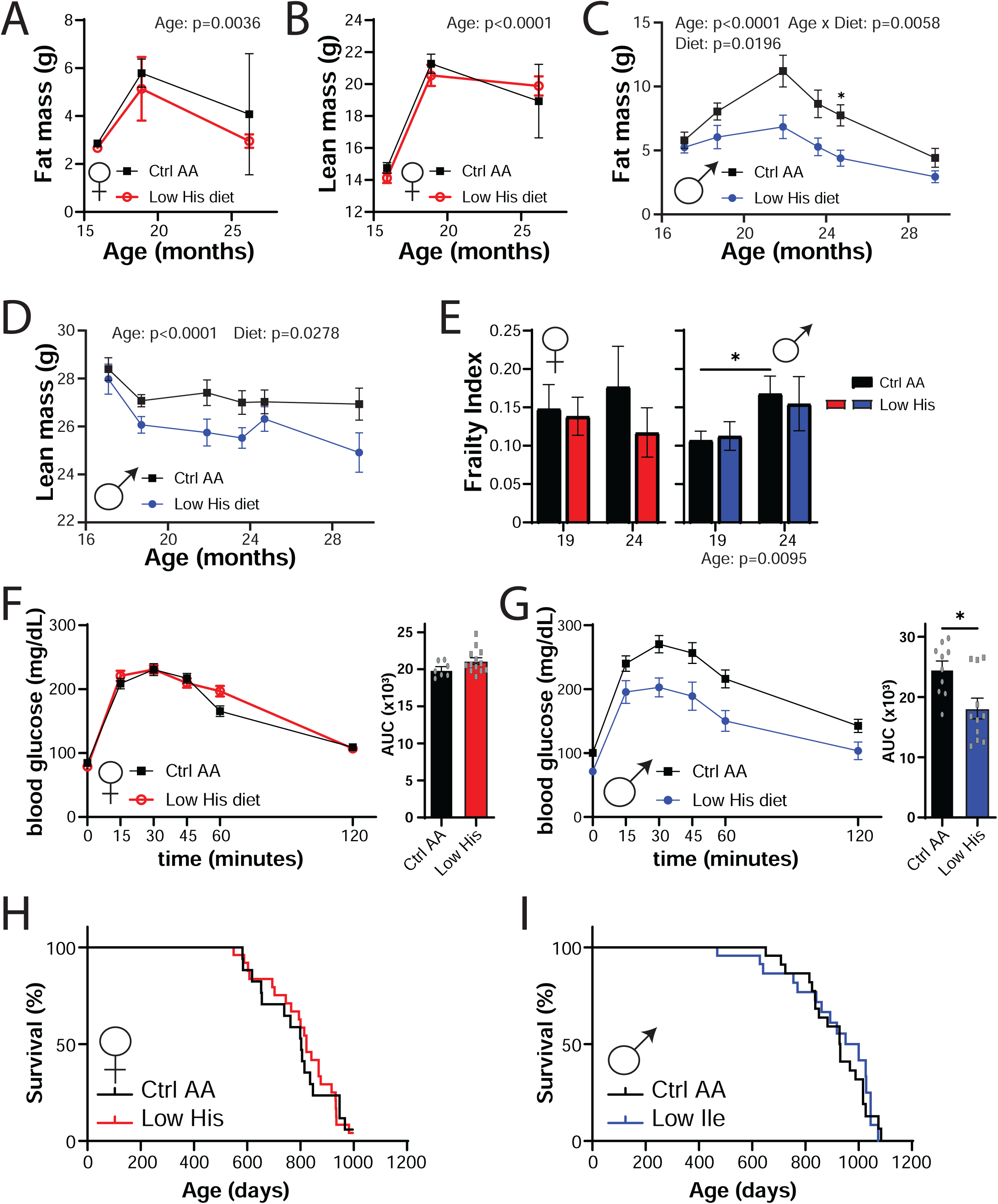
Mid-life histidine improves metabolic health in males but does not impact lifespan. Male and female C57BL/6J.Nia mice were placed on Control AA and Low His diets starting at 16 months of age. (A-D) Fat mass and lean mass were determined at multiple time points (n= n varies by month; maximum n = 9 Ctrl AA females, 10 Low His females, 14 Ctrl AA males, 10 Low His males). (E) Frailty was assessed at 19 and 24 months of age. (A-E) Statistics for the overall effects of diet, age and the interaction represent the P value from a mixed-effects model (restricted maximum likelihood (REML)), *p<0.05, two-sided Sidak’s post hoc test. (F-G) Glucose tolerance was determined in (F) 23 month old females and (G) 25 month old males (n = 7-13/group females and 10-11/group males, *p < 0.05, t-test). (H-I) Kaplan–Meier plots showing the survival of female (H) and male (I) C57BL/6J.Nia mice fed the indicated diets starting at 16 months of age.

Both mice and humans become increasingly frail with age, and we utilized a recently developed mouse frailty index to assess the effect of a Low His diet frailty (Whitehead *et al*., 2014; Kane *et al*., 2016). In male mice we observed an overall effect of age on frailty, but in neither sex did a Low His diet result in a significant change in frailty (**Fig. 10E**). We also assessed glucose tolerance as the mice aged, and in 25-month-old male mice, we found that a Low His diet improved glucose tolerance (**Figs. 10F-10G**). Despite the improvements in metabolic health, a Low His diet had no significant effect on the overall survival of either male or female mice when started at 16 months of age (**Figs. 10H-10I**).

### Decreased dietary histidine protects from the effects of a Western diet

We next decided to examine whether decreasing dietary histidine can protect from the deleterious effects of a Western diet (**Fig. 11A**). We fed young 6-week-old C57BL/6J mice either an amino acid-defined WD or a WD Low His diet as described above; in parallel a group of mice was fed a Control AA-defined diet. Over the next 12 weeks, we found that mice fed the WD Low His diet were completely protected from the effects of a WD diet on weight and fat mass, while experiencing a transient minor dip in lean mass (**Fig. 11B-11C**). WD-fed mice had significantly thicker layer of dermal WAT at the end of 12 weeks than Control AA-fed mice; the WD Low His mice did not show this increase (**Fig. 11D**). The overall effect was a complete protection from the effect of a WD on adiposity (**Fig. 11E**).

**Figure 11.**
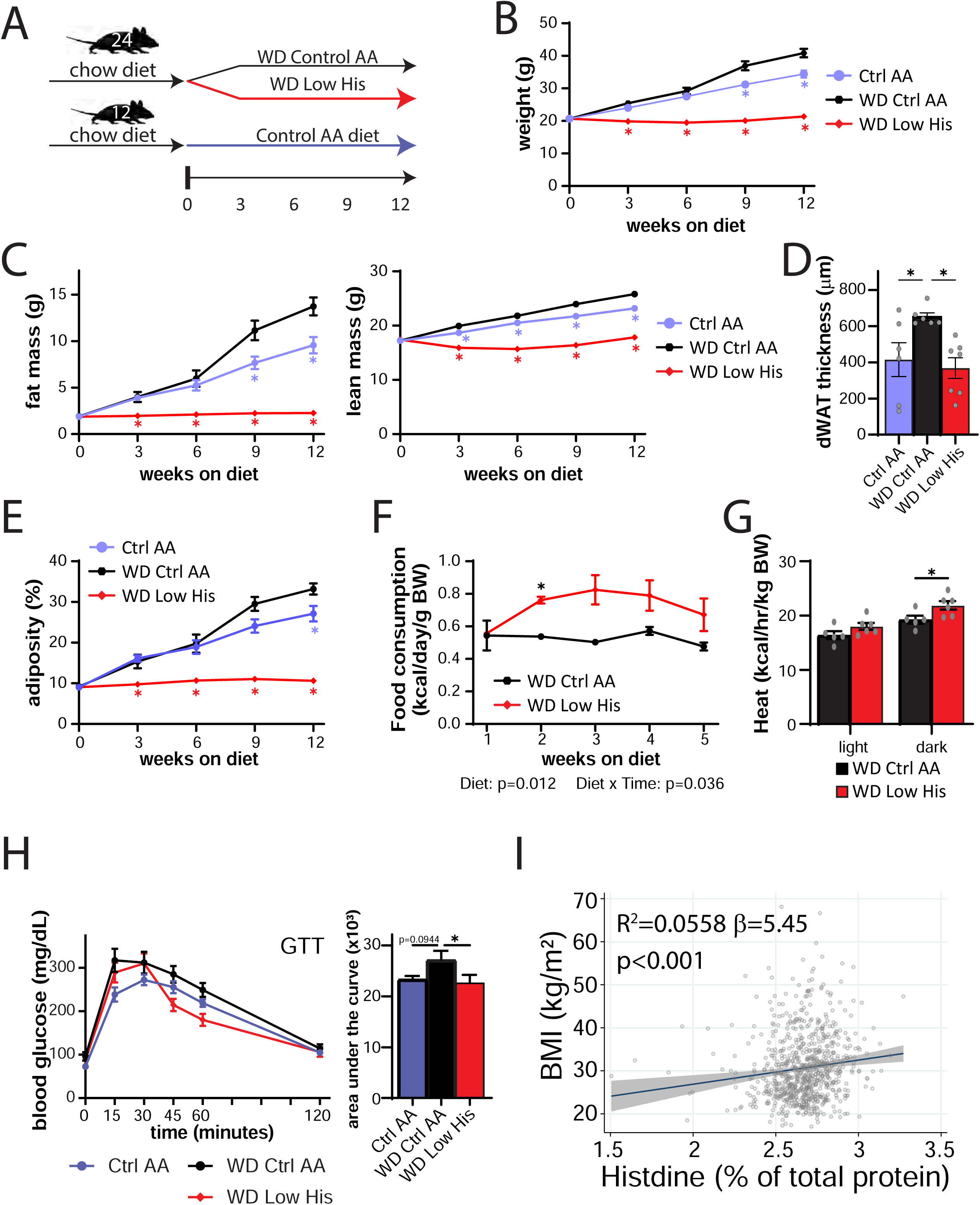
Decreased histidine intake prevents diet-induced obesity and glucose intolerance, and is associated with lower BMI in humans. (A) Experimental scheme. (B-E) Body weight (B), fat mass and lean mass (C), dermal WAT thickness (D), and body composition (E) of mice consuming the indicated diets. (B, C, E) n=6-12 mice per group, *p<0.05 vs. all other groups, Tukey’s test following a mixed-effects model (Restricted Maximum Likelihood (REML). (D) n=6-7/group, *p<0.05 vs. WD Ctrl AA, Dunnett’s test post ANOVA. (F) Food consumption (n=6 cages per group, *p<0.05, Sidak’s test post 2-way RM ANOVA). (G) Energy expenditure (heat) normalized to body weight of mice fed the indicated diets (n=5-6 mice per group, *p<0.05, Sidak’s test post 2-way RM ANOVA). (H) Glucose tolerance test after 3 weeks on the indicated diets (n=12/group, *p<0.05 vs. WD Ctrl AA, Dunnett’s test post ANOVA). (I) Association between BMI and percent of total protein from histidine from the SHOW study (n = 788, shaded area represents 95% CI). Data represented as mean ± SEM.

The effect of the WD Low His diet on weight was not driven by reduced calorie intake; indeed as we anticipated based on our results above, WD Low His-fed mice actually consumed more calories relative to WD Ctrl AA-fed mice (**Fig. 11F**). Instead, mice fed a WD Low His diet have increased energy expenditure (**Fig. 11G**). WD Low His-fed mice also had improved glucose tolerance relative to WD Ctrl AA-fed mice (**Fig. 11H**).

### Increased histidine intake is associated with higher BMI in humans

As reducing dietary histidine has potent beneficial effects on the body composition of mice, we next examined whether dietary levels of histidine are associated with human health. Body mass index (BMI) is easily measured and highly correlated with fat mass in humans (Peltz *et al*., 2010; Ranasinghe *et al*., 2013; Bell *et al*., 2018). Among a randomly selected, cross-sectional population-based sample of adults (>18 years old), we examined associations between estimated histidine intake relative to total protein and BMI (kg/m^2^) determined from measured weight and height. After adjusting for confounding variables, we discovered that a single percentage point increase in dietary histidine intake relative to total protein—for example, from 2% to 3% of protein—is linked with a 5.45 (95% CI: 2.4-8.5) unit rise in BMI (p<0.001) **(Fig. 11I).** The effects were similar and remained highly significant when males and females were considered separately, and adding an interaction term to the regression model was non-significant. In accordance with our findings in animals, our data show that a greater dietary consumption of protein that contains a larger proportion of histidine is associated with an increased BMI in both males and females in a general population-based sample. These findings, when seen in the context of our animal investigations, show that dietary protein quality – specifically, the proportion of the diet may also be a key regulator of human weight, adiposity, and metabolic health.

## Discussion

Recent studies have demonstrated that protein quality – in particular, the specific amino acid composition of the diet – plays a significant role in mediating the effects of dietary protein on health. Much of the initial research in this area focused on the essential amino acid methionine, restriction of which promotes metabolic health and extends lifespan in rodents (Orentreich *et al*., 1993; Richie *et al*., 1994; Miller *et al*., 2005; Lees *et al*., 2014; Schmidt *et al*., 2016; Lees *et al*., 2017), due to the belief that lower levels of dietary methionine might account for the health benefits of a vegan diet (McCarty *et al*., 2009; MacArthur *et al*., 2021). The BCAAs, leucine, isoleucine, and valine, have also been the subject of significant attention, as blood levels of BCAAs in humans are associated with diabetes and insulin resistance (Newgard *et al*., 2009). Our lab has previously demonstrated that restriction of the three dietary BCAAs improves metabolic health, fitness, and extends the lifespan of mice (Fontana *et al*., 2016; Cummings *et al*., 2018; Richardson *et al*., 2021; Yu *et al*., 2021). We have shown that restriction of the BCAA isoleucine is necessary and sufficient for the metabolic benefits of a low protein diet; further, in humans dietary isoleucine levels are associated with increased body mass index, while the blood level of isoleucine in humans is associated with mortality (Deelen *et al*., 2019; Yu *et al*., 2021).

Our previous experiments with BCAAs suggested that one or more of the other essential AAs – histidine, lysine, methionine, phenylalanine, threonine, and tryptophan – contributed to the effects of a low protein diet on body composition. In this work, we analyzed the effect of restricting each of these single essential AAs by 67% relative to the Control diet. We observed statistically significant effects of restricting either histidine, phenylalanine, or threonine on fat mass accretion and adiposity, and restriction of either histidine or phenylalanine on body weight. While the overall effect of restricting any of these three amino acids was one of reduced adiposity, mice restricted in either histidine or phenylalanine also gained less lean mass during the study. The possible consequences of reduced lean mass gain for metabolic health is an area for future investigation, and a limitation of the present study is the lack of analysis of growth-associated hormones, including growth hormone and IGF-1, that might potentially mediate the effects of these AAs on fat and lean mass.

We concentrated our follow-up analysis in diet induced obese mice on phenylalanine and histidine restriction due to the larger effect size of these interventions, as well as the recent exploration of dietary threonine restriction by another group (Yap *et al*., 2020). Here, we determined that specifically reducing dietary histidine, without altering energy density, the caloric contribution of AAs to the diet, or the protein:carbohydrate ratio (Solon-Biet *et al*., 2014) is sufficient to restore metabolic health. To the best of our knowledge, these studies represent the first examination of a reduced histidine diet in a mouse model of pre-existing diet-induced obesity and type 2 diabetes. Specifically reducing histidine rapidly normalizes the weight of diet-induced obese mice without calorie restriction, even as these mice continue to eat an otherwise high-fat, high-sucrose Western diet. In contrast, the effects of phenylalanine restriction on body composition were observed, with a significant reduction of body weight in WD Low Phe-fed mice.

While the effects of histidine restriction on body composition are similar to the effects of restricting BCAAs, they are distinct in that restriction of dietary histidine in lean mice has no effects on glucose tolerance. Further, while in obese mice histidine restriction modestly improves glucose homeostasis, unlike restriction of isoleucine, histidine restriction does not improve hepatic insulin sensitivity; instead, the benefits of histidine restriction on glucose tolerance and insulin sensitivity closely tracks the changes in weight and decreased adiposity. This was surprising as a WD Low His diet has dramatic and beneficial effects on hepatic steatosis, which we would have expected to improve hepatic insulin sensitivity. The molecular mechanism by which histidine restriction reduces liver fat is also unclear; while we found significant effect on the expression of a number of liver genes involved with lipogenesis, similar effects were observed in WD Low AA- fed mice, which did not show a reduction in liver fat.

The decreased weight and adiposity of histidine-restricted animals was not associated not with decreased caloric intake, but instead with increased calorie intake and increased energy expenditure. In this respect histidine restriction is similar to the effects of protein restriction, methionine restriction, BCAA and isoleucine restriction, as well as threonine and tryptophan restriction, in C57BL/6J male mice (Laeger *et al*., 2014; Hill *et al*., 2017; Wanders *et al*., 2017; Cummings *et al*., 2018; Yap *et al*., 2020; Richardson *et al*., 2021; Green *et al*., 2022; Hill *et al*., 2022). Restriction of protein, BCAAs/isoleucine, and methionine in C57BL/6J male mice activate the FGF21-UCP1 axis, and these effect of these restriction on food intake and energy expenditure require *Fgf21* (Laeger *et al*., 2014; Hill *et al*., 2017; Wanders *et al*., 2017; Yu *et al*., 2021).

We were therefore surprised to find that while histidine restriction increases food consumption, energy expenditure, and increased insulin-stimulated glucose uptake into iWAT and BAT, histidine restriction did not induce hepatic *Fgf21* or iWAT *Ucp1*. As FGF21 can be produced by many tissues other than the liver, we tested the role of FGF21 directly using *Fgf21^-/-^* mice, we found that indeed the effects of histidine restriction do not depend on the presence of *Fgf21*; male *Fgf21^-/-^* mice show the same effects on body composition in response to histidine restriction that wild-type males do, as well as similar (if perhaps blunted) effects on food consumption and energy expenditure. Thus, in combination these results indicate that the metabolic effects of histidine restriction are partially or wholly independent of the FGF21-UCP1 axis.

Additional research will be required to fully define the physiological and molecular mechanisms by which dietary histidine regulates energy balance and body composition. One possibility is that histidine restriction regulates energy balance not via thermogenesis, but via changes in the regulation of heat loss through the skin. Key modulators of heat rate loss via the skin include insulation by dermal WAT (dWAT) (Alexander *et al*., 2015; Kasza *et al*., 2019), and we observe that mice fed a WD Low His diet have significantly thinner dWAT than WD Ctrl AA- fed controls. This could potentially explain the slightly lower core body temperature of WD Low His-fed mice, as well as the cold-resistance of mice fed a WD Low His mice, as mice initially subject to a slight cold stress may be well-equipped to activate compensatory mechanisms to stabilize body temperature. More research will be required to examine this possibility and understand if and how histidine restriction regulates heat loss.

Another possibility is suggested by the fact that histidine is a precursor for the formation of histamine, which regulates allergies and inflammatory responses, and alpha ketoglutarate (AKG). Histamine has been shown to act as an anorexigenic agent via the regulation of leptin (Itateyama *et al*., 2003; Yoshimatsu, 2008), and dietary histidine has been shown to suppress food intake in rats (Kasaoka *et al*., 2004). These results, as well as recent work showing that genetic inhibition of histamine synthesis has been shown to protect from hepatic steatosis (Kennedy *et al*., 2018), are consistent with our results that restriction of dietary histidine increases food consumption and has beneficial effects on hepatic steatosis. Histidine is a precursor to AKG, and we would therefore expect histidine restriction to lower AKG levels; however as AKG promotes beneficial metabolic effects, we would expect this to negatively, not positively, impact metabolic health. Future work should examine how mice distribute limited supplies of histidine to protein synthesis and catabolic needs, and how the levels of histamine and AKG, as well as other metabolites, are impacted by changes in dietary histidine levels.

Interestingly, we found that restriction of methionine, lysine, or tryptophan had only minor effects on metabolic health. The lack of an effect of methionine restriction (MR) was the most surprising, as multiple studies have shown methionine restriction extends the lifespan of rodents and improves metabolic health, improving glucose tolerance and reducing adiposity (Orentreich *et al*., 1993; Miller *et al*., 2005; Lees *et al*., 2014). However, the degree of MR in these studies is typically 80% or more, substantially more than the level of restriction we examined; similarly, recent work on tryptophan restriction has utilized a great degree of restriction and a lower absolute level of protein (Yap *et al*., 2020). Further, recent work has shown that the presence of dietary cysteine – which in mammals is synthesized from methionine – blocks the metabolic effects of MR (Wanders *et al*., 2016). This effect of cysteine likely results from a combination of decreased MR-induced oxidative stress, and a decreased demand for methionine-derived cysteine. In our current study, we have not directly measured the levels of AAs or their metabolites in blood or tissue, and thus the negligible impact of methionine, lysine, or tryptophan restriction in our study may result from our intervention not significantly changing blood or tissue levels of these AAs. Finally, as we analyzed the effects of methionine, lysine, or tryptophan only in young lean male mice, it is possible that these amino acids would play a role in regulating body composition or metabolic health in obese, aged, or female mice.

We also examined the effect of histidine restriction as a midlife intervention to promote healthy aging and longevity. Interestingly, we observed a sexually dimorphic effect of histidine restriction on body composition, with histidine restriction marginally reducing fat mass without impacting lean mass in aged females. In contrast, aged male mice fed a Low His diet lost both fat mass and lean mass. While lean mass loss could be detrimental, especially in aged individuals, we did not observe a negative effect on either frailty or lifespan. Glucose tolerance also improved selectively in males and not females. Future studies will need to carefully assess if this effect on lean mass in aged mice is detrimental to healthspan though not reflected in our frailty assessments.

Other limitations of our study include that our analysis of the effects of histidine restriction on young and diet-induced obese mice were conducted exclusively in male mice of a single inbred strain. Over the last few years, it has become apparent that both sex and genetic background play an important role in the response to dietary interventions (Mitchell *et al*., 2016; Green & Lamming, 2021; Roy *et al*., 2021); our own previous work has shown that protein or BCAA restriction has distinct metabolic and molecular effects when examined in both sexes and different strains of mice (Richardson *et al*., 2021; Green *et al*., 2022). In our analysis of histidine restriction as a midlife intervention, we observed distinct effects of a Low His diet on body composition and glucose tolerance in males and females. Future studies should consider the effect of dietary histidine levels on the metabolic health of multiple genetic backgrounds and both sexes. Finally, our findings in humans are only correlative, and it remains to be determined if and how dietary histidine regulates body composition in humans. Notably, histidine is not the only amino acid that we have found to be associated with BMI – we previously reported that restriction of isoleucine (but not leucine or valine) is also associated with BMI to a lesser degree (Yu *et al*., 2021). Determining the contribution of other essential and non-essential amino acids to BMI is an important area for further research.

In conclusion, we have found that dietary histidine is a critical modulator of metabolic health and body composition in mice and potentially in humans (**Figure 8I**). Specifically reducing dietary histidine reduces weight and fat mass accretion in young lean mice, and rapidly normalizes body weight, fat mass, and adiposity in diet-induced obese mice without requiring calorie restriction. These metabolic effects likely result from increased energy expenditure. In contrast to other dietary regimens involving the restriction of dietary protein or specific AAs, restriction of histidine does not increase energy expenditure through the FGF21-UCP1 axis.

Further, we find that reduction of dietary histidine in diet-induced obese mice also corrects metabolic abnormalities associated with obesity, including impaired glucose tolerance, impaired insulin sensitivity, and hepatic steatosis. Restriction of dietary histidine also protects young lean mice from Western diet-induced obesity, weight gain, and glucose intolerance. While dietary histidine restriction promotes metabolic health in aged male mice, it does not extend their lifespan. Finally, we find that dietary levels of histidine are correlated with metabolic health in humans, and we linked dietary levels of histidine to higher BMI. Our results highlight the critical importance of dietary histidine levels for obesity and metabolic health of both mice and humans, and suggest that reducing dietary histidine levels may be a novel therapeutic and public health strategy to combat the obesity epidemic.

## Materials and Methods

All procedures were performed in conformance with institutional guidelines and were approved by the Institutional Animal Care and Use Committee of the William S. Middleton Memorial Veterans Hospital, institutional assurance number D16-00403 (Madison, WI, USA) or the University of Pennsylvania, institutional assurance number D16-00045. (Philadelphia, PA, USA). The research complies with the policies of *The Journal of Physiology* and the animal ethics checklist (Grundy, 2015).

### Mouse information

Except as noted otherwise, all the studies described here were conducted at the the William S. Middleton Memorial Veterans Hospital (Madison, WI, USA) using male C57BL/6J mice purchased from The Jackson Laboratory (Bar Harbor, ME, USA), and all mice were acclimated to the animal research facility for at least one week before entering studies.

To test the effects of individual amino acid restriction, 10-week-old male mice were placed on control or individual amino acid restricted diets as indicated. To test the metabolic effects of histidine in the context of diet-induced obesity, 6-week-old male C57BL/6J mice were pre- conditioned with WD (TD.88137, Envigo, Madison, WI) for 12 weeks, and another group of mice were fed control chow diet (Purina 5001). Following 12-weeks of WD preconditioning, mice on WD diet were then switched to the amino acid defined Western diets described in Figures, while mice fed a chow diet were switched to an amino acid defined Control diet.

To generate male FGF21 KO mice, we crossed CMV-Cre mice (Schwenk *et al*., 1995) from the Jackson Laboratory (006054) with mice expressing a floxed allele of *FGF21* (Potthoff *et al*., 2009) from The Jackson Laboratory (022361), then crossed with C57BL/6J mice to remove CMV-Cre; genotyping was performed as described (Yu *et al*., 2021). Briefly, PCR was performed using DreamTaq DNA Polymerase (Thermo Scientific, EP0705) according to insert instructions.

All amino acid defined diets were obtained from Envigo, and diet compositions and item numbers are provided in **Tables 1 and 2**. Mice were housed in a specific pathogen free (SPF) mouse facility with a 12:12 hour light/dark cycle and with free access to food and water, and were maintained at a temperature of approximately 22°C, except for the cold stress study described below. Animals were group housed in static microisolator cages, except when temporarily housed in a Columbus Instruments Oxymax/CLAMS metabolic chamber system. Group sizes are provided in the figure legends.

### *In vivo* Procedures

For glucose tolerance tests, food was withheld overnight (for 16 hours), and mice were then injected intraperitoneally with glucose (1 g/kg) as described (Yu *et al*., 2021). Insulin tolerance tests were performed by fasting mice for 16 hours (Figures S1 and S2) or 4 hours (Figure S3) starting at lights on, and then injecting insulin (0.75U/kg) intraperitoneally. Glucose measurements were taken using a Bayer Contour blood glucose meter and test strips. Body composition was assessed using an EchoMRI Body Composition Analyzer (EchoMRI, Houston, TX, USA) (Bellantuono *et al*., 2020). To assess metabolic physiology (O2, CO2, RER, and food consumption) and spontaneous activity, mice were placed into a Columbus Instruments Oxymax/CLAMS or CLAMS-HC metabolic chamber system (Columbus Instruments, Columbus, OH, USA) and acclimated for approximately 24 hours prior to data collection, and data from a continuous 24 hour period was then selected for analysis (Bellantuono *et al*., 2020).

### Hyperinsulinemic-Euglycemic Clamp and Temperature Studies

All hyperinsulinemic-euglycemic clamp and temperature studies were performed in mice at the Penn Diabetes Research Center Rodent Metabolic Phenotyping Core (University of Pennsylvania); mice were sourced directly from The Jackson Laboratory and diets were obtained from Envigo (Madison, WI). Mice were preconditioned with Western diet (TD.88137) and then switched to either the WD Ctrl AA diet or WD Low His diet.

5 days after the diet switch, rectal temperature was taken at 22°C. Mice were then placed in a cage pre-equilibrated to 4°C for one hour. Upon removal from the 4°C cage, rectal temperature was taken again.

Hyperinsulinemic-euglycemic clamp studies of obese, Western diet fed animals were performed as follows. Indwelling jugular vein and carotid artery catheters were surgically implanted in the mice for infusion 7 days prior to the clamp study day as previously described (Ayala *et al*., 2006). Mice were fasted for 5 hours prior to initiation of clamp and acclimated to the containers (plastic bowl with alpha dry). Jugular vein and arterial line are connected to the dual swivel 2 hours prior to the clamp initiation. A [3-^3^H] glucose infusion was primed (1.5 μCi) and continuously infused for a 90 min equilibration period (0.075 μCi/min). Baseline measurements were determined in arterial blood samples collected at -10 and 0 min (relative to the start of the clamp) for analysis of glucose, [3-^3^H] glucose specific activity, basal insulin and free fatty acids. The clamp was started at t = 0 min with a primed-continuous infusion of human insulin (2.5 mU/Kg/min; Novolin Regular Insulin), a donor blood infusion at 4.5 uL/min to prevent a 5% drop in the hematocrit, and glucose (D50 mixed with [3-^3^H] glucose 0.05 µCi/uL) was infused at variable glucose infusion rate (GIR) to maintain euglycemia. The mixing of D50 with [3-^3^H] glucose is required to maintain the specific activity constant during the clamp period. Arterial blood samples were taken at t=80-120 min for the measurement of [3-^3^H] glucose specific activity, clamped insulin and free fatty acid levels. 120 minutes after initiation of clamp, ^14^C-2DG (12µCi) is injected and arterial blood samples obtained at 2, 5, 10, 15 and 25 min to determine Rg, an index of tissue-specific glucose uptake in various tissues. After the final blood sample, animals were injected with a bolus of pentobarbital, and tissues were collected and frozen in liquid nitrogen and stored at -80°C for subsequent analysis.

Processing of samples and calculations: Radioactivity of [3-^3^H] glucose, [^14^C]2DG and [^14^C]2DG- 6-phosphate were determined as previously described (Ayala *et al*., 2007). The glucose turnover rate (total Ra; mg/kg/min) was calculated as the rate of tracer infusion (dpm/min) divided by the corrected plasma glucose specific activity (dpm/mg) per kg body weight of the mouse. Glucose appearance (Ra) and disappearance (Rd) rates were determined using steady-state equations and endogenous glucose production (Ra) was determined by subtracting the GIR from total Ra. Tissue specific glucose disposal (Rg; µmol/100g tissue/min) was calculated as previously described (Ayala *et al*., 2007). The plasma insulin concentration was determined by mouse ELISA kit (Biomarker core, University of Pennsylvania).

### Assays and Kits

Unless noted otherwise, all assays were performed using samples from overnight 16-hour fasted mice. Blood FGF21 levels were assayed by a mouse/rat FGF-21 quantikine ELISA kit (MF2100) from R&D Systems (Minneapolis, MN, USA).

### Histology and Quantitative PCR

Tissues were harvested from mice on the indicated diets following overnight (16-hr) fasting. A portion of the liver was embedded in OCT, and then cryosectioned and Oil-Red-O stained by the UWCCC Experimental Pathology Laboratory. Liver sections were imaged using an EVOS microscope as previously described (Cummings *et al*., 2018). For quantification of lipid droplet size in liver, 6 independent fields were obtained for tissue from each mouse by investigators blinded to the treatment group and then quantified using NIH ImageJ. Skin was paraformaldehyde-fixed (4%) overnight, and then paraffin-embedded for evaluation (Kasza *et al*., 2014).

A portion of the liver and iWAT was harvested and snap frozen in liquid nitrogen for subsequent molecular analysis. For quantitative PCR, RNA was extracted from liver or adipose tissue using Tri-reagent (Sigma-Aldrich, St. Louis, MO, USA). 1 mg of RNA was used to generate cDNA using Superscript III (Invitrogen, Carlsbad, CA, USA). Oligo dT primers and primers for real-time PCR were obtained from Integrated DNA Technologies (Coralville, IA, USA). Reactions were run on an Applied Biosystems StepOne Plus machine (Applied Biosystems, Foster City, CA, USA) with Sybr Green PCR Master Mix (Invitrogen, Carlsbad, CA, USA). Actin was used to normalize the results from gene-specific reactions.

### SHOW study

We analyzed the association between dietary histidine intake and body mass index (BMI) in a sample of 2016-2017 Survey of Health of Wisconsin (SHOW) participants. SHOW is an ongoing population-based health examination survey of non-institutionalized residents of Wisconsin. Detailed survey methods have been previously described (Nieto *et al*., 2010; Malecki *et al*., 2022). Survey components relevant to the current analysis included an in-home interview accompanied by measurements of weight and height, and a self-administered questionnaire including the National Cancer Institute’s Diet History Questionnaire from which specific dietary intake variables were derived.

The study population included 788 individuals that completed all parts of the survey including diet history, blood draw and exam visit. Demographic characteristics of the population are described in Table 3. All study protocols were approved by the University of Wisconsin Health Sciences Institutional Review Board, and all participants provided written informed consent as part of the initial home visit. The intake of histidine was estimated from the Diet History Questionnaire II (National Cancer Institute) using Diet*Calc software (National Cancer Institute). The estimated levels of histidine are expressed as the percent (%) of total protein. The primary outcome was measured body mass index (kg/m2) derived from measured weight (kg) and height (m). Multiple linear regression was performed using STATA 17.0 (STATA Corp LLC, College Station, TX) with BMI as the outcome and percent of total protein from histidine as the predictor of interest. In the model we also adjusted for age, gender, education, income, total caloric intake, and physical activity. Linear regression models were stratified by gender to mirror mouse model findings and interactions including gender and percent histidine were considered.

**Table 3.** SHOW Demographics. SHOW population demographics.

| **SHOW Demographics** | **Estimate**<br>**N=788** | **95% Confidence Interval (Logit)** |  |
| --- | --- | --- | --- |
| Age, mean (SD) | 53.5 (16.3) | 52.4 | 54.7 |
| Male (%) | 42.1 | 38.7 | 45.6 |
| Body Mass Index<br>(kg/m2), mean (SD) | 30.6 (7.7) | 17.0 | 68.1 |
| Body Mass Index<br>(kg/m2), categories (%) |  |  |  |
| < 25 | 25.1 | 22.2 | 28.3 |
| 25<30 | 28.6 | 25.5 | 31.8 |
| >= 30 | 46.3 | 42.8 | 49.8 |
| Education (%) |  |  |  |
| <=High School | 6.5 | 5.1 | 8.6 |
| High School Diploma or<br>General Education<br>Degree (GED) | 38.9 | 35.6 | 42.4 |
| >High school | 54.4 | 50.9 | 57.9 |
| Race Ethnicity (%) |  |  |  |
| Non Hispanic White | 79.7 | 76.7 | 82.3 |
| Non-Hispanic Black<br>(alone or in<br>combination) | 12.8 | 10.7 | 15.4 |
| Hispanic (any race) | 3.8 | 2.7 | 5.4 |
| Other (non-Hispanic) | 3.7 | 2.6 | 5.3 |

### Quantification and statistical analysis

Unless otherwise stated, statistical analysis was conducted using Prism 7/8(GraphPad Software). Tests involving repeated measurements were analyzed with two-way repeated-measures ANOVA, followed by a Tukey-Kramer or Dunnett’s post-hoc test as specified. All other comparisons of three or more means were analyzed by one-way ANOVA followed by a Dunnett’s or Tukey-Kramer post-hoc test as specified in the figure legends. Additional comparisons, if any, were corrected for multiple comparisons using the Bonferroni method.

### Data availability

Source data for all figures is available in Supporting Information, with the exception of sensitive human data. Human data are available for qualified investigators upon request from Dr. Kristen Malecki and the Survey of Health of Wisconsin. A fee for service model may apply for access to restricted data.

## Author contributions

VF, ABS, NER, DY, EPC, JMR, IK, CLG, CMA, JAB, KCM, and DWL contributed to the conception or design of the work. VF, ABS, MS, NER, DY, GES, MET, RB, EPC, JMR, SEY, MHW, RH, IK, JLT, CLG, CD, CMA, JAB, KCM, and DWL contributed to data acquisition, analysis or interpretation. All authors contributed to drafting or revision of the manuscript, and approved the final version. All authors agree to be accountable for all aspects of the work in ensuring that questions related to the accuracy or integrity of any part of the work are appropriately investigated and resolved. All authors agree that all persons designated as authors qualify for authorship, and all those who qualify for authorship are listed.

## Acknowledgements

We thank Tina Herfel (Envigo) for help with diet design, Maria Nikodemova for assistance with accessing SHOW data, and Susanne Keipert for helpful discussions.

