## Supplemental Figures and Table S3 for "Regulation of metabolic health by dietary histidine in mice" (from another version of this preprint: version 1, 2022-04-24)

**Supplemental Material**

#### Supplemental Figure Legends

**Figure S1. Dietary restriction of histidine, phenylalanine or threonine alters energy balance.**

(A-D) Metabolic chambers were utilized to assess multiple components of energy balance, including food consumption (A), energy expenditure (heat) (B), spontaneous activity (C), and respiratory exchange ratio (RER) (D); n=3-6 per group, *$p<0.05$, Dunnet's test vs. WD Ctrl AA conducted separately for the light and dark cycles post 2-way RM ANOVA. (E) Insulin tolerance test (n=8-9/group, *$p<0.05$ vs. Ctrl AA, Dunnett's test post ANOVA). Data represented as mean ± SEM.

**Figure S2. Dietary restriction of lysine, methionine, or tryptophan has no effect on energy balance or insulin sensitivity.**

(A-D) Metabolic chambers were utilized to assess multiple components of energy balance, including food consumption (A), energy expenditure (heat) (B), spontaneous activity (C), and respiratory exchange ratio (RER) (D); n=8-9 per group, *$p<0.05$, Dunnet's test vs. WD Ctrl AA conducted separately for the light and dark cycles post 2-way RM ANOVA. (E) Insulin tolerance test (n=9/group, *$p<0.05$ vs. Ctrl AA, Dunnett's test post ANOVA). Data represented as mean ± SEM.

**Figure S3. Restricting dietary histidine alters energy balance and insulin sensitivity of diet-induced obese mice.**

(A-D) Metabolic chambers were utilized to assess multiple components of energy balance, including food consumption (A), energy expenditure (heat) (B), spontaneous activity (C), and respiratory exchange ratio (RER) (D); n=5-8 per group, *$p<0.05$, Dunnet's test vs. WD Ctrl AA conducted separately for the light and dark cycles post 2-way RM ANOVA. (E) Insulin tolerance test (n=9-12/group, *$p<0.05$ vs. Ctrl AA, Dunnett's test post ANOVA). Data represented as mean ± SEM.

#### Supplemental Table Legends

**Table S1. Diet composition of diets used to investigate the effect of reduced levels of essential amino acids in the context of a normal calorie diet.**

**Table S2. Diet composition of diets used to investigate the effect of reduced levels of essential amino acids in the context of a WD.**

**Table S3. SHOW population demographics.**

### Figure S1

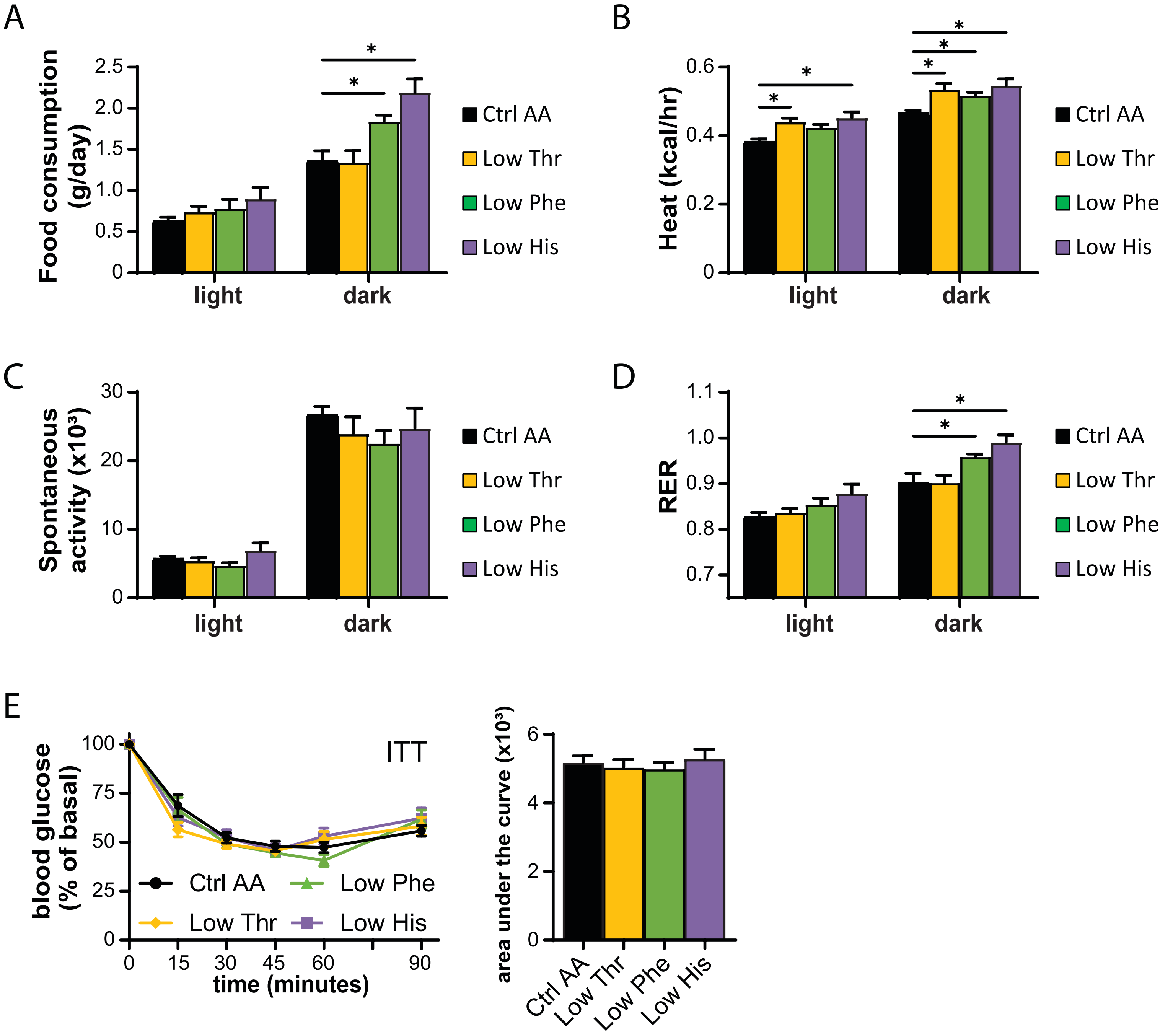

A
Food consumption (g/day)
2.5
2.0
1.5
1.0
0.5
0
light
dark
*
*
Ctrl AA
Low Thr
Low Phe
Low His
B
Heat (kcal/hr)
0.6
0.4
0.2
0
light
dark
*
*
*
*
*
Ctrl AA
Low Thr
Low Phe
Low His
C
Spontaneous activity (x10³)
30
20
10
0
light
dark
Ctrl AA
Low Thr
Low Phe
Low His
D
RER
1.1
1.0
0.9
0.8
0.7
light
dark
*
*
Ctrl AA
Low Thr
Low Phe
Low His
E
blood glucose (% of basal)
100
75
50
25
0
ITT
Ctrl AA
Low Phe
Low Thr
Low His
0
15
30
45
60
90
time (minutes)
area under the curve (x10³)
6
4
2
0
Ctrl AA
Low Thr
Low Phe
Low His

### Figure S2

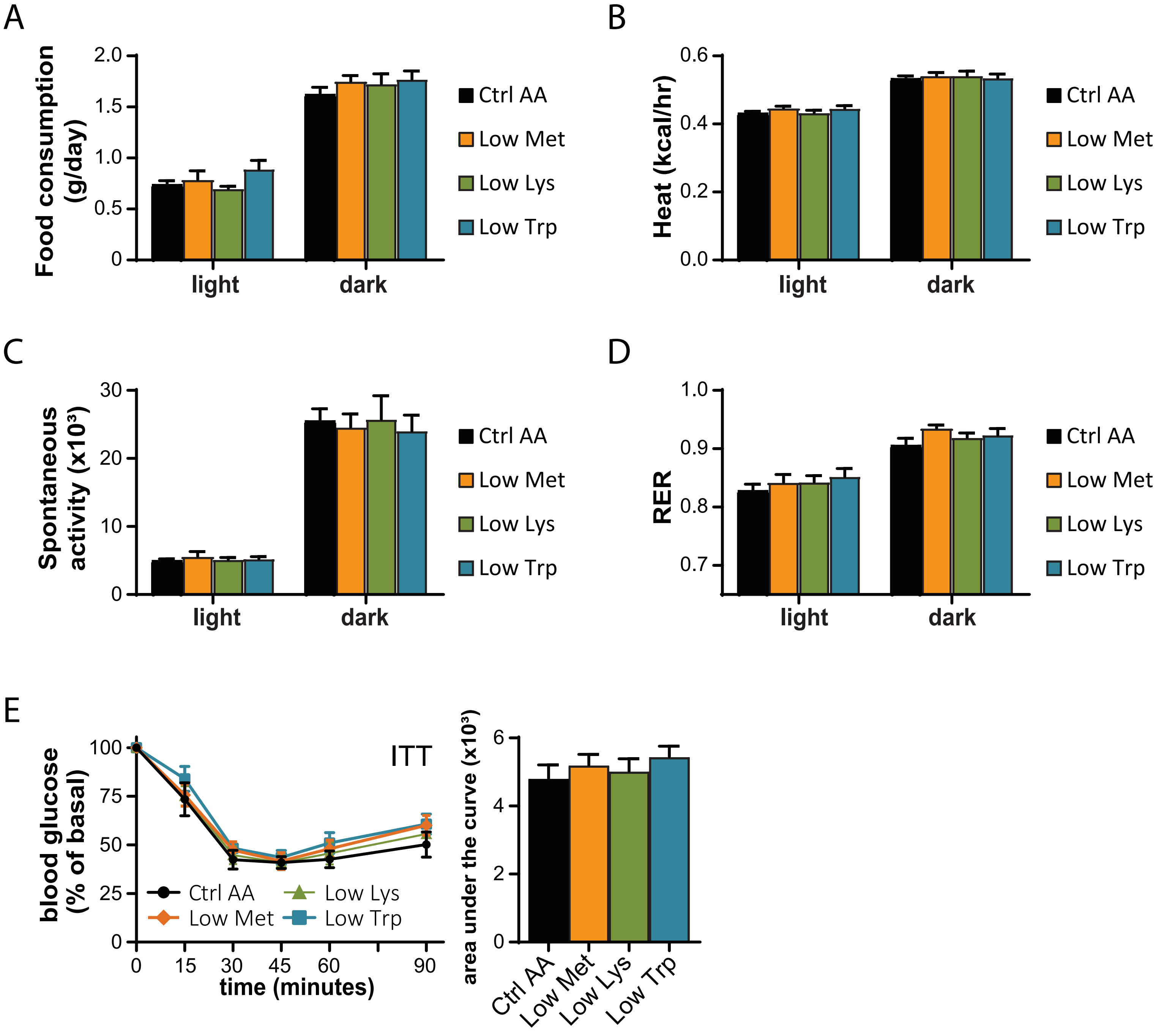
A
Food consumption (g/day)
2.0
1.5
1.0
0.5
0
light
dark
Ctrl AA
Low Met
Low Lys
Low Trp
B
Heat (kcal/hr)
0.6
0.4
0.2
0.0
light
dark
Ctrl AA
Low Met
Low Lys
Low Trp
C
Spontaneous activity (x10³)
30
20
10
0
light
dark
Ctrl AA
Low Met
Low Lys
Low Trp
D
RER
1.0
0.9
0.8
0.7
light
dark
Ctrl AA
Low Met
Low Lys
Low Trp
E
blood glucose (% of basal)
100
75
50
25
0
ITT
Ctrl AA
Low Lys
Low Met
Low Trp
0
15
30
45
60
90
time (minutes)
area under the curve (x10³)
6
4
2
0
Ctrl AA
Low Met
Low Lys
Low Trp

### Figure S3

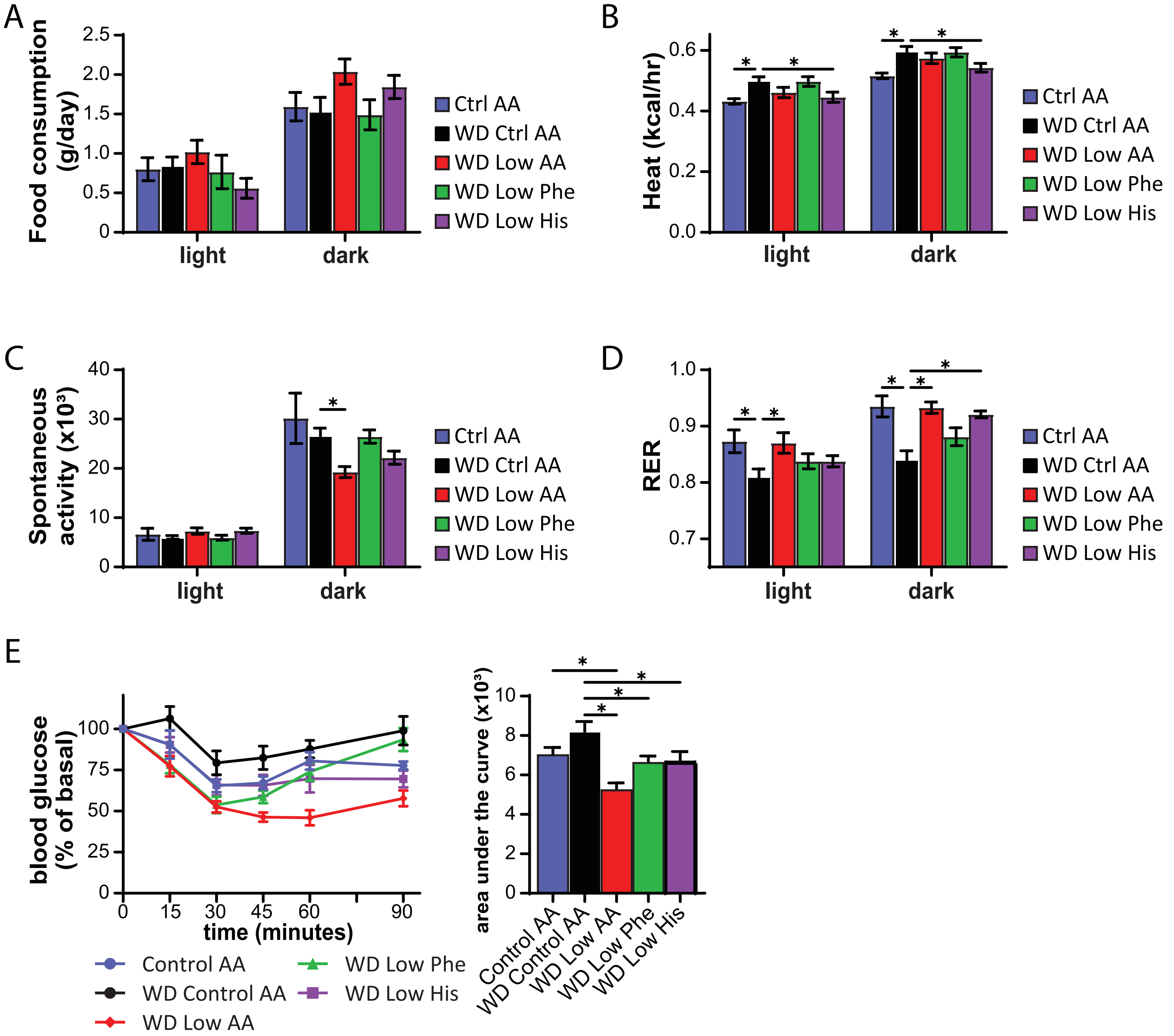

A
Food consumption (g/day)
2.5
2.0
1.5
1.0
0.5
0
light
dark
Ctrl AA
WD Ctrl AA
WD Low AA
WD Low Phe
WD Low His
B
Heat (kcal/hr)
0.6
0.4
0.2
0.0
light
dark
Ctrl AA
WD Ctrl AA
WD Low AA
WD Low Phe
WD Low His
C
Spontaneous activity (x10³)
40
30
20
10
0
light
dark
Ctrl AA
WD Ctrl AA
WD Low AA
WD Low Phe
WD Low His
D
RER
1.0
0.9
0.8
0.7
light
dark
Ctrl AA
WD Ctrl AA
WD Low AA
WD Low Phe
WD Low His
E
blood glucose (% of basal)
100
75
50
25
0
0
15
30
45
60
90
time (minutes)
Control AA
WD Control AA
WD Low AA
WD Low Phe
WD Low His
area under the curve (x10³)
10
8
6
4
2
0
Control AA
WD Control AA
WD Low AA
WD Low Phe
WD Low His

Table S3: SHOW population demographics

| SHOW Demographics | |
|---|---|
| **Age, mean ( SD)** | 53.6 (16.4) |
| **male, n (%)** | 333 (42.2) |
| **BMI, mean (SD)** | 30.6 (7.7) |
| **BMI categories:** | |
| <25, n (%) | 189 (25.1) |
| 25<30, n (%) | 226 (28.6) |
| ≥ 30, n (%) | 365 (46.2) |
| **Education:** | |
| < high school, n (%) | 53 (6.72) |
| HS Diploma/GED, n (%) | 307 (38.9) |
| > high school, n (%) | 429 (54.4) |
| **Race** | |
| Non-Hispanic White, n (%) | 628 (79.7) |
| Black/African American, n (%) | 101 (12.8) |
| Hispanic, n (%) | 30 (3.8) |
| Other, n (%) | 29 (3.7) |
